# Multi-modality neuroimaging brain-age in UK Biobank: relationship to biomedical, lifestyle and cognitive factors

**DOI:** 10.1101/812982

**Authors:** James H Cole

## Abstract

The brain-age paradigm is proving increasingly useful for exploring ageing-related disease and can predict important future health outcomes. Most brain-age research utilises structural neuroimaging to index brain volume. However, ageing affects multiple aspects of brain structure and function, which can be examined using multi-modality neuroimaging. Using UK Biobank, brain-age was modelled in n=2,205 healthy people with T1-weighted MRI, T2-FLAIR, T2*, diffusion-MRI, task fMRI and resting-state fMRI. In a held-out healthy validation set (n=520), chronological age was accurately predicted (r=0.79, mean absolute error=3.52 years) using LASSO regression, higher than using any modality separately. Thirty-four neuroimaging phenotypes were deemed informative by the regression (after bootstrapping); predominantly grey-matter volume and white-matter microstructure measures. When applied to new individuals from UK Biobank (n=14,701), significant associations with multi-modality brain-predicted age difference (brain-PAD) were found for: stroke history, diabetes diagnosis, smoking, alcohol intake and some, but not all, cognitive measures (corrected p<0.05). Multi-modality neuroimaging can improve brain-age prediction, and derived brain-PAD values are sensitive to biomedical and lifestyle factors that negatively impact brain and cognitive health.

## 1. Introduction

Ageing has a pronounced effect on the human brain, resulting in cognitive decline and an increased risk of neurodegenerative diseases and dementia. Though brain ageing is ubiquitous, differences between individuals can be substantial. Some people experience cognitive decline, neurodegeneration and age-related brain diseases in midlife, while others retain the majority of their cognitive function well into their tenth decade.

Poorer brain-health during ageing has a pronounced negative impact for individuals, their families and for society (Wasay, et al., 2016). Hence, ways of identifying people at risk of poorer brain health during ageing have become an important research goal. One such approach, the so-called ‘brain-age paradigm’ (Cole and Franke, 2017, Franke and Gaser, 2019), aims to define the brain’s ‘biological age’ (Jackson, et al., 2003, Ludwig and Smoke, 1980). The idea here is that genetic and environmental influences can influence the rate at which age-associated biological changes accumulate, and that one’s biological age might be a better predictor of disease-risk, functional capacity and residual lifespan, than chronological age. By defining a statistical model of healthy brain ageing, using neuroimaging data to predict chronological age, one can then evaluate neuroimaging data from new individuals, and consider whether their ‘brain-age’ appears younger or older than their chronological age.

Having an older-appearing brain has previously associated with markers of physiological ageing (e.g., grip strength, lung function, walking speed) and cognitive ageing (Cole, et al., 2018), suggesting that brain and body ageing are related (Cole, et al., 2019c). An older brain age has also been associated with poor future outcomes, including progression from mild cognitive impairment to dementia (Franke and Gaser, 2012, Gaser, et al., 2013) and mortality (Cole, et al., 2018). Brain-age can also be moderated by a range of different neurological and psychiatric diseases (reviewed in Cole, et al., 2019b).

Statistical methods for modelling brain age using neuroimaging are generally highly accurate, with most approaches able to account for >90% of the variance in chronological age. The mean absolute error (MAE) of predictions are generally around 4-5 years (when predicting between 18-90 years), though the winners of a recent competition achieved MAE < 3 years using an ensemble deep-learning method (https://www.photon-ai.com/pac2019). Nevertheless, further improving model accuracy is important step towards application in clinical settings, where predictions will be made at the individual level. Potential clinical uses include screening for poorer brain-health in cognitively normal middle-aged adults, in stratifying clinical trial recruitment or as a surrogate outcome measure of neuroprotective treatments.

Most brain-age models use only T1-weighted structural MRI, reflecting brain volumes. Given that ageing impacts many aspects of brain structure and function, and that these can be measured with other neuroimaging modalities, one could potentially improve accuracy by incorporating complementary data on brain connectivity, white-matter hyperintensities, iron deposition and brain activity during tasks and at rest. A multi-modality approach has the benefit of providing a richer and more comprehensive explanation of the mechanisms underlying individual differences in brain ageing. For example, changes in white-matter microstructure may precede alterations in brain volume, or there may be regional differences in the susceptibility to age-related brain changes. Including more measurements should capture more age-related variance and thus improve model accuracy. Multi-modality imaging could also be used to generate an array of modality-specific brain “ages”. For example, an individual could have a structural brain-age, a diffusion brain-age and a functional connectivity brain-age. In healthy people, these separate “ages” should be closely related, according to the concept of biological age, but for patient groups, distinct patterns of aberrant brain ageing could emerge.

Several previous brain-age studies have used two or three modalities (Cherubini, et al., 2016, Groves, et al., 2012, Liem, et al., 2017, Richard, et al., 2018). Liem and colleagues (2017) studied n=2354 people, finding improved accuracy when combining T1-weighted structural MRI with resting-state fMRI (MAE = 4.29 years). Structural MRI resulted in higher accuracy than fMRI when used independently. Meanwhile, Groves and colleagues (2012) using linked independent component analysis to merge T1-weighted and diffusion-MRI in n=484 participants, finding that the resulting largest component could predict age with high accuracy (correlation between chronological age and brain age *r* = 0.95). Similarly, Richard and colleagues (2018) used T1-weighted and diffusion-MRI in their study of n=877 participants. Again, the combination of modalities achieved top performance (*r* = 0.86, MAE = 6.14 years), while T1-derived values resulted in higher age-prediction accuracy than white-matter microstructure measures from diffusion-MRI. Cherubini and colleagues (2016) used three modalities; T1-weighted, T2* relaxometry and diffusion-MRI data. This small study (n=140) achieved excellent performance (*r* = 0.96) with these three modalities, although accuracy was still generally high with single modalities.

Findings from these multi-modality brain age studies suggest that T1-weighted MRI is the best single predictor, but that adding other modalities results in higher accuracy than in any single modality. These results are promising, but still limited to two or three modalities. This is largely due to data availability; the UK Biobank imaging study (Miller, et al., 2016) presents a new opportunity to overcome this limitation. With a planned total of n=100,000 individuals undergoing standardised neuroimaging, using four identical, dedicated MRI scanners, a wealth of multi-modality MRI data are becoming available. The modalities include T1-weigthed MRI, T2-FLAIR, susceptibility-weighted imaging (SWI), diffusion-MRI, task fMRI and resting-state fMRI. In addition to neuroimaging, participants provide detailed information on current health, lifestyle and medical history, as well as participating in cognitive testing and providing blood for biological assessments.

Here, I tested whether brain-age prediction in healthy people can be improved by combining data from these six modalities, as well as evaluating their independent performance. I then applied the resulting multi-modality brain-age model to a held-out sample of UK Biobank participants to test the relationship between brain-prediction age and factors relating to health, lifestyle and cognitive function. I hypothesised that i) highest accuracy (i.e., lowest MAE, highest variance explained in age) would be achieved by combining data from all modalities ; ii) that T1-weighted MRI would provide the highest independent accuracy; iii) that all modalities could be used to significantly predict chronological age independently; iv) that smoking, alcohol intake, major physiological health conditions and poorer cognition would be associated with having an older-appearing brain.

## 2. Materials and Methods

### 2.1. Participants

Data from n=22,392 participants from UK Biobank were analysed. Of these, n=20,237 completed the brain MRI assessment; which included n=2,776 participants who were missing one or more neuroimaging modality. This left n=17,461 participants (aged 45-80 years, n=9,274 females, n=8,187 males) with complete neuroimaging data. Two subsets were then defined; one for training the statistical model (the training set), one for testing associations with other UK Biobank measures (the test set). Training data included only people who met criteria for being healthy at the time of scanning. Exclusion criteria were an ICD-10 diagnosis, a self-reported long-standing illness disability or infirmity (UK Biobank data field #2188), no self-reported diabetes (field #2443), no stroke history (field #4056), not having good or excellent self-reported health (field #2178). This gave n=2,725 (mean age = 61.47 ± 7.2 years, 1,343 females and 1,382 males). The test set comprised the remaining n=14,701 participants (mean age = 62.64 ± 7.5, 7,914 females, 6,787 males). All UK Biobank data were downloaded and reformatted using the R package ukbtools (Hanscombe, et al., 2019).

All participants provided informed consent. UK Biobank has ethical approval from the North West Multi-Centre Research Ethics Committee (MREC). Further details on the UK Biobank Ethics and Governance framework are here: https://www.ukbiobank.ac.uk/the-ethics-and-governance-council/. The present analyses were conducted under data application number 40933, *Optimising neuroimaging biomarkers of brain ageing to identify genetic and environmental risk factors for poor brain health*.

### 2.2. Data acquisition

Full details on the UK Biobank neuroimaging data are provided here: https://biobank.ctsu.ox.ac.uk/crystal/crystal/docs/brain_mri.pdf. In brief, T1-weighted MRI used an MPRAGE sequence with 1mm isotropic resolution. The T2 protocol uses a fluid-attenuated inversion recovery (FLAIR) contrast with the 3D SPACE optimized readout, with a 1.05×1×1mm resolution. The SWI protocol used a dual-echo 3D gradient echo acquisition at 0.8×0.8×3mm resolution, (TEs = 9.4, 20ms). T2* values (i.e., signal decay times) were estimated from the magnitude images at the two echo times. Diffusion-MRI data were acquired with two b-values (b = 1,000, 2,000 s/mm_2_) at 2mm isotropic resolution, with a multiband acceleration factor of 3. For both diffusion-weighted shells, 50 diffusion-encoding directions were acquired (covering 100 distinct directions over the two b-values). Task and resting-state fMRI use the following acquisition parameters, with 2.4-mm spatial resolution and TR = 0.735s, with a multiband acceleration factor of 8. The total neuroimaging acquisition protocol lasted 32 minutes per participant.

### 2.3. Data processing

Data used in the current analysis were the imaging-derived phenotypes developed centrally by researchers involved in UK Biobank (Miller, et al., 2016) and distributed via the data showcase (http://biobank.ctsu.ox.ac.uk/crystal/index.cgi). These data were the available summary metrics for T1-weighted MRI, diffusion-MRI, T2-FLAIR, SWI (i.e., T2*) and task fMRI. Data from resting state fMRI was so-called bulk data, in this case the 25-dimension partial correlation matrices (data field #25752). These matrices had been converted into vectors of length 210, representing the pairwise correlations between the BOLD timeseries from 21 separate resting state networks (details on page 16 here: https://biobank.ctsu.ox.ac.uk/crystal/crystal/docs/brain_mri.pdf). Four for 25 networks were classified as noise by UK Biobank researchers and removed prior to being made available. The figure of 210 values is arrived at by 21 networks multiplied by 20 (i.e., excluding the identity correlation), then divided by 2 (i.e., as the matrix is diagonally symmetrical). The use of partial correlation aims to provide a better estimate of direct ‘connection’ strengths between networks than using full correlations.

The final set of neuroimaging phenotypes numbered as follows: T1-weighted = 165, T2-FLAIR = 1 (total volume of white-matter hyperintensities), T2* = 14, diffusion-MRI = 657, task fMRI = 14, resting-state fMRI = 210. This gave 1,079 phenotypes in total (see Table A1 in Appendix).

### 2.4. Statistical analysis

All statistical analyses were carried using R version 3.5.2 (R Core Team, 2015). The R Markdown notebook containing all statistical code can be found here: https://james-cole.github.io/UKBiobank-Brain-Age/.

To define a multi-modality healthy ageing model, the training data were randomly split into separate training (80%, n=2,205) and validation sets (20%, n=520), to ensure that model accuracy could be evaluated in an unbiased manner. All neuroimaging phenotypes were normalised (i.e., scaled by standard deviation and mean centred) to account for the different measurement scales used by the different modalities.

To predict age from neuroimaging data, a least absolute shrinkage and selection operator (LASSO) regression was run, with age as the outcome variable and neuroimaging phenotypes as the predictors. LASSO regression imposes an L1-norm penalty, in which the goal is to minimise the absolute value of the beta coefficients in the model. Coefficients that shrink below a threshold (lambda) are automatically set to zero. This leads to a sparse solution, which reduces the variance in the model (i.e., regularisation); in effect working to select informative features and remove uninformative ones. To derive the optimal value lambda, ten-fold cross-validation was first run using a range of lambdas, and the highest value within one standard error of the minimum was used in subsequent analysis.

Since LASSO can be susceptible to biased solutions caused by multicollinearity, bootstrapping (i.e., resampling with replacement) was used to generate a distribution of coefficients for each predictor variable. Here, 1000 bootstraps were run, and the mean coefficient and 95% confidence interval (calculated using the basic bootstrap method) were computed. Neuroimaging phenotypes whose bootstrapped 95% confidence intervals did not overlap zero were considered to be informative for the prediction of age.

The LASSO regression procedure was then run per modality (T1-weighted, T2-FLAIR, diffusion-MRI, SWI, task fMRI, resting-state fMRI), where phenotypes from that modality alone were included. This time bootstrapping was not conducted, as the goal was to compare model-level performance across modalities, rather than identify important features within each model.

To test for associations between brain-age and health-related variables in UK Biobank, the LASSO model was applied to predict age in people from the testing set. As recent research has highlighted a proportional bias in brain-age calculation, whereby the difference between chronological age and brain-predicted age is negatively correlated with chronological age (Le, et al., 2018, Liang, et al., 2019, Smith, et al., 2019), an age-bias correction procedure was employed. This entailed calculating the regression line between age and brain-predicted age in the training set, then using the slope (i.e., coefficient) and intercept of that line to adjust brain-predicted age values in the testing set (by subtracting the intercept and then dividing by the slope). After applying age-bias correction the brain-predicted age difference (brain-PAD) was calculated; chronological age subtracted from brain-predicted age. This gives a resulting value in unit years, with positive values representing an older-appearing brain and negative values a younger-appearing brain.

Associations between brain-PAD values and demographic, biomedical, cognitive and lifestyle measures were then tested, using linear regression models. In the models, brain-PAD was the outcome measure, the variable of interest was a predictor, alongside age, age_2_, sex, height, volumetric scaling from T1-weighted MRI to standard (data field #25000) and mean task fMRI head motion (averaged across space and time points; data field #25742) as covariates. These covariates were chosen due to face validity (i.e., theoretically likely to relate to brain structure) and statistical correlation with brain-PAD.

Biomedical measures (from visit #2, the imaging visit) tested were systolic blood pressure, diastolic blood pressure, weight, body mass index [BMI], hip circumference, diabetes diagnosis; stroke diagnosis, and facial ageing. Lifestyle measures were smoking status alcohol intake frequency, duration of moderate activity or vigorous activity per day. Cognitive performance measures were fluid intelligence score, Trail making task: duration to complete numeric path trail 1, duration to complete alphanumeric path trail 2, Matrix pattern completion: number of puzzles correctly solved, duration spent answering each puzzle and Tower rearranging: number of puzzles correct. Full details on the coding for these variables are reported in the UK Biobank data showcase (http://biobank.ctsu.ox.ac.uk/crystal/search.cgi). Of particular note, the coding for alcohol intake frequency has lower values reflecting higher intake (1 = Daily or almost daily; 2 = Three or four times a week; 3 = Once or twice a week; 4 = One to three times a month; 5 = Special occasions only; 6 = Never; −3 = Prefer not to answer). Multiple testing correction for these 18 separate measures was conducted using false-discovery rate correction (Benjamini and Hochberg, 1995).

## 3. Results

### 3.1. Demographics of healthy participants in training and test set

After subdividing UK Biobank into a healthy training set and separate test set, the two groups were broadly comparable (Table 1). Age was slightly higher in the test, which also had a higher proportion of females. The groups were equivocal in terms of blood pressure, BMI, weight and hip circumference. By design, rates of stroke history and a diagnosis of diabetes were higher in the test set (1.4% and 5.7% respectively), as these were exclusion criteria for the healthy training set.

**Table 1.** UK Biobank neuroimaging participant characteristics.

|  | Healthy training set | Test set |
| --- | --- | --- |
| N | 2725 | 14701 |
| Age, mean ± SD | 61.47 ± 7.21 | 62.64 ± 7.45 |
| Female, % (n) | 49.3% (1343) | 53.8% (7914) |
| Body mass index, median [IQR] | 25.94 [23.58-28.68] | 25.94 [23.56-28.88] |
| Weight, kg, median [IQR] | 75 [65.6-85.4] | 74.6 [65.2-85] |
| Hip circumference, cm, median [IQR] | 100 [95.5-105] | 100 [96-106] |
| Diastolic blood pressure, median [IQR] | 79 [72-87] | 78 [71-85] |
| Systolic blood pressure, median [IQR] | 139 [126-151.75] | 137 [125-150] |
| ICD-10 diagnosis, % (n) | 0% (0) | 73% (12753) |
| Diabetes, % (n) | 0% (0) | 5.72% (836) |
| Stroke, % (n) | 0% (0) | 1.37% (201) |

### 3.2. Multi-modality neuroimaging can predict chronological age in healthy people

When applying the brain-model model to the validation dataset, the correlation between chronological age and brain-predicted age was r=0.786, R_2_=0.618, MAE=3.515, with an age-bias of r=−0.65 (Figure 1).

**Figure 1.**
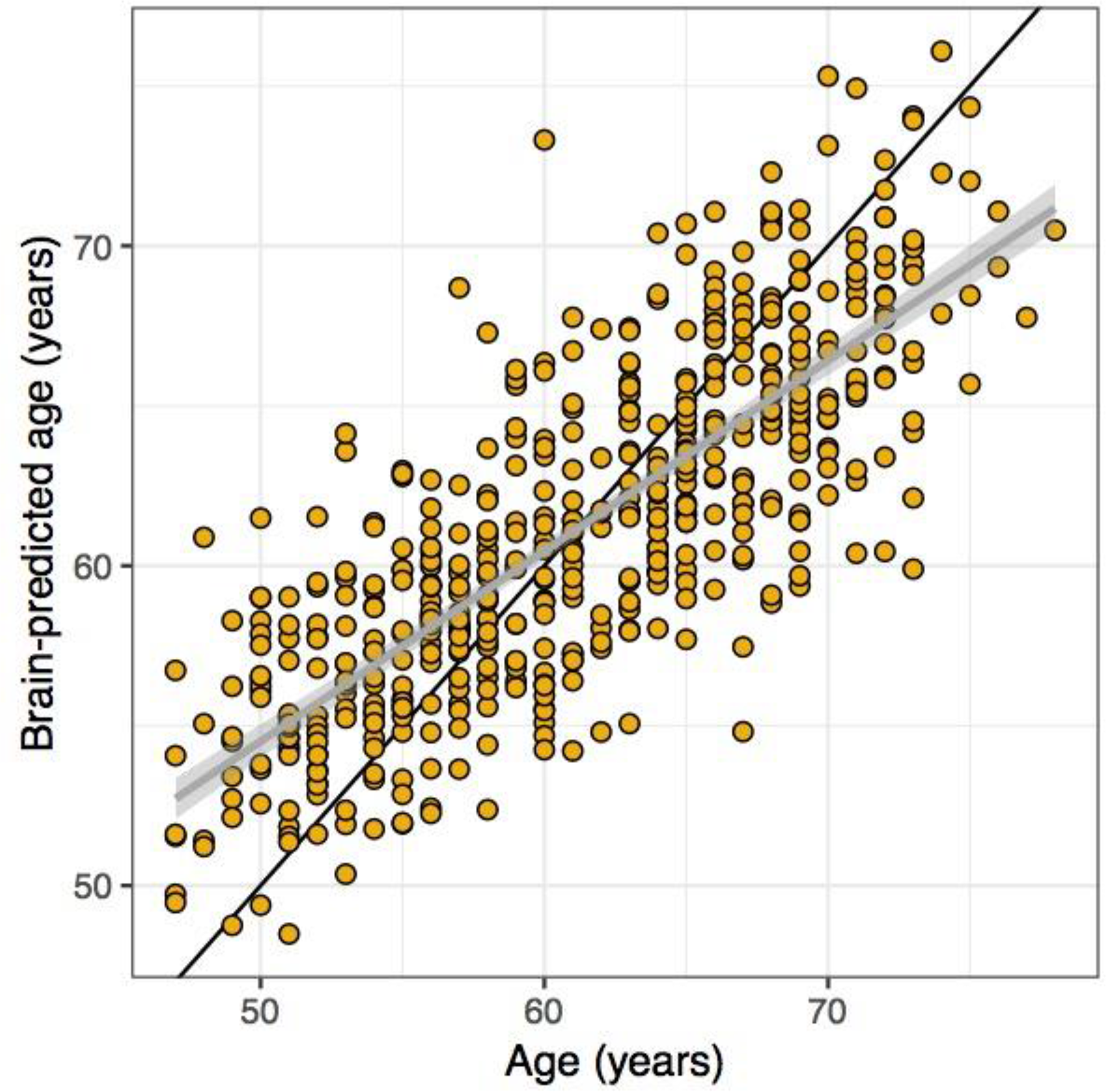
Brain-predicted age from multi-modality LASSO regression model. Scatterplot depicting chronological age (x-axis) by brain-predicted age (y-axis) in UK Biobank validation set (n=520). Black line is the line of identity. Grey line is the regression line of age on brain-predicted age with shaded errors representing the 95% confidence intervals.

From the initial input of 1,079 neuroimaging phenotypes, 221 were set to non-zero in the LASSO regression model. After bootstrapping, 34 neuroimaging phenotypes had 95% confidence intervals that did not overlap zero (Table 2). This is despite 886 out of the 1,079 neuroimaging phenotypes being significantly correlated with age at p < 0.05; even using Bonferroni correction 704 neuroimaging were significantly correlated with age (adjusted p = 4.6*10_−5_).

**Table 2.** Informative neuroimaging phenotypes consistently for predicting age in a multi-modality LASSO regression.

| Neuroimaging phenotype | UK Biobank data field # | Modality | Coefficient [95% confidence intervals] |
| --- | --- | --- | --- |
| Volume of grey matter normalised for head size | 25005 | T1-weighted | -1.466 [-1.851, -1.242] |
| Weighted mean ICVF in tract forceps minor | 25661 | Diffusion-MRI | -1.029 [-1.789, -0.699] |
| Volume of brain stem 4th ventricle | 25025 | T1-weighted | 0.692 [0.484, 1.139] |
| Volume of grey matter in ventral striatum left | 25890 | T1-weighted | -0.666 [-1.022, -0.393] |
| Weighted mean L1 in tract anterior thalamic radiation right | 25572 | Diffusion-MRI | 0.513 [0.218, 1.026] |
| Mean ISOVF in fornix on FA skeleton | 25445 | Diffusion-MRI | 0.501 [0.219, 1.003] |
| Mean FA in middle cerebellar peduncle on FA skeleton | 25056 | Diffusion-MRI | -0.477 [-0.91, -0.275] |
| Volume of grey matter in putamen right | 25883 | T1-weighted | 0.460 [0.262, 0.911] |
| Volume of thalamus right | 25012 | T1-weighted | -0.454 [-0.909, -0.116] |
| Mean FA in cerebral peduncle on FA skeleton left | 25071 | Diffusion-MRI | -0.453 [-0.906, -0.226] |
| Mean FA in superior cerebellar peduncle on FA skeleton left | 25069 | Diffusion-MRI | 0.429 [0.053, 0.859] |
| Mean L1 in middle cerebellar peduncle on FA skeleton | 25200 | Diffusion-MRI | -0.428 [-0.815, -0.162] |
| Mean L3 in posterior thalamic radiation on FA skeleton right | 25324 | Diffusion-MRI | 0.418 [0.129, 0.835] |
| Mean L2 in fornix cres/stria terminalis on FA skeleton left | 25287 | Diffusion-MRI | 0.393 [0.062, 0.787] |
| Mean ICVF in body of corpus callosum on FA skeleton | 25347 | Diffusion-MRI | 0.381 [0.031, 0.762] |
| Mean L1 in anterior limb of internal capsule on FA skeleton left | 25217 | Diffusion-MRI | 0.349 [0.095, 0.698] |
| Mean MO in fornix cres/stria terminalis on FA skeleton left | 25191 | Diffusion-MRI | -0.336 [-0.61, -0.061] |
| Volume of grey matter in vi cerebellum right | 25899 | T1-weighted | -0.330 [-0.661, -0.059] |
| Volume of grey matter in frontal operculum cortex right | 25863 | T1-weighted | -0.321 [-0.582, -0.146] |
| Mean OD in superior longitudinal fasciculus on FA skeleton left | 25433 | Diffusion-MRI | 0.316 [0.169, 0.633] |
| Volume of grey matter in vermis crus ii cerebellum | 25904 | T1-weighted | 0.306 [0.115, 0.532] |
| Weighted mean L3 in tract uncinate fasciculus left | 25648 | Diffusion-MRI | 0.302 [0.002, 0.603] |
| Volume of putamen left | 25015 | T1-weighted | -0.284 [-0.567, -0.011] |
| Mean MD in cingulum hippocampus on FA skeleton right | 25140 | Diffusion-MRI | -0.272 [-0.544, -0.109] |
| Mean L2 in splenium of corpus callosum on FA skeleton | 25252 | Diffusion-MRI | -0.271 [-0.542, -0.014] |
| Weighted mean MO in tract anterior thalamic radiation left | 25544 | Diffusion-MRI | 0.263 [0.065, 0.526] |
| 90th percentile of BOLD effect in group-defined mask for shapes activation | 25761 | Task fMRI | 0.229 [0.05, 0.425] |
| Weighted mean MO in tract forceps minor | 25553 | Diffusion-MRI | -0.226 [-0.452, -0.031] |
| Median T2* in putamen left | 25030 | T2* | -0.221 [-0.442, -0.016] |
| Volume of grey matter in thalamus right | 25879 | T1-weighted | 0.217 [0.005, 0.435] |
| Median BOLD effect in group-defined mask for faces-shapes contrast | 25048 | Task fMRI | -0.212 [-0.423, -0.06] |
| Weighted mean L1 in tract parahippocampal part of cingulum left | 25575 | Diffusion-MRI | -0.174 [-0.348, -0.001] |
| Resting-state partial correlation 25-dimension IC v157 | 25752 | Resting-state fMRI | 0.165 [0.024, 0.329] |
| Mean L2 in cingulum cingulate gyrus on FA skeleton right | 25282 | Diffusion-MRI | -0.119 [-0.237, -0.009] |

ICVF = intracellular volume fraction; FA = fractional anisotropy; ISOVF = isotropic volume fraction; MO = mode of anisotropy; OD = orientation dispersion; v# = variable from resting-state fMRI partial correlation matrix, using 25-dimension independent component analysis.

These 34 ‘informative’ neuroimaging phenotypes were predominantly from T1-weighted MRI or diffusion-MRI, although T2*, task fMRI and resting-fMRI variables were included. Given that the variables were scaled across modalities, the relative beta coefficients from the model can be considered. The variable with the largest absolute coefficient was the volume of grey matter (normalised for head size) beta = −1.466, 95% confidence intervals [−1.85, −1.24], suggesting that lower grey matter volume is associated with higher brain-age estimates. Lower weighted mean intra-cellular volume fraction (ICVF) in the forceps minor was also associated with higher brain-age estimates, suggesting that neurite density in the anterior corpus callosum and related regions is lower in older-appearing brains. Increasing volume of the fourth ventricle, equating to an enlargement of this CSF space, was also associated with higher brain-age estimates. Other notable associations include reductions in volume and changes in diffusion metrics in the cerebellum.

### 3.3. Single modality brain-age prediction

Variables from each neuroimaging modality were used separately to train a model to predict age in the validation dataset (Table 3). The best performing modalities (i.e., highest correlation with age, lowest MAE) were T1-weighted MRI and diffusion-MRI. The other four modalities were only able to explain a limited amount of variance in age, particularly the task fMRI. The pattern of correlations across different “ages” predicted by each modality is shown in Figure 2. The highest correlation between two “ages” is between T1-weighted and diffusion-MRI (*r* = 0.73).

**Table 3.** Brain-age prediction performance from individual modalities.

| Modality | Number of variables | Correlation between age and brain-age ($r$) | Variance in age explained ($R^2$) | Mean Absolute Error (years) | Age bias (correlation between age and age-difference) |
| --- | --- | --- | --- | --- | --- |
| T1-weighted | 165 | 0.685 | 0.470 | 4.132 | -0.716 |
| T2-FLAIR | 1 | 0.308 | 0.095 | 5.653 | -0.942 |
| T2* | 14 | 0.325 | 0.105 | 5.739 | -0.981 |
| Diffusion-MRI | 675 | 0.668 | 0.447 | 4.282 | -0.815 |
| Task fMRI | 14 | 0.164 | 0.027 | 5.920 | -0.984 |
| Resting-state fMRI | 210 | 0.434 | 0.189 | 5.302 | -0.933 |

**Figure 2.**
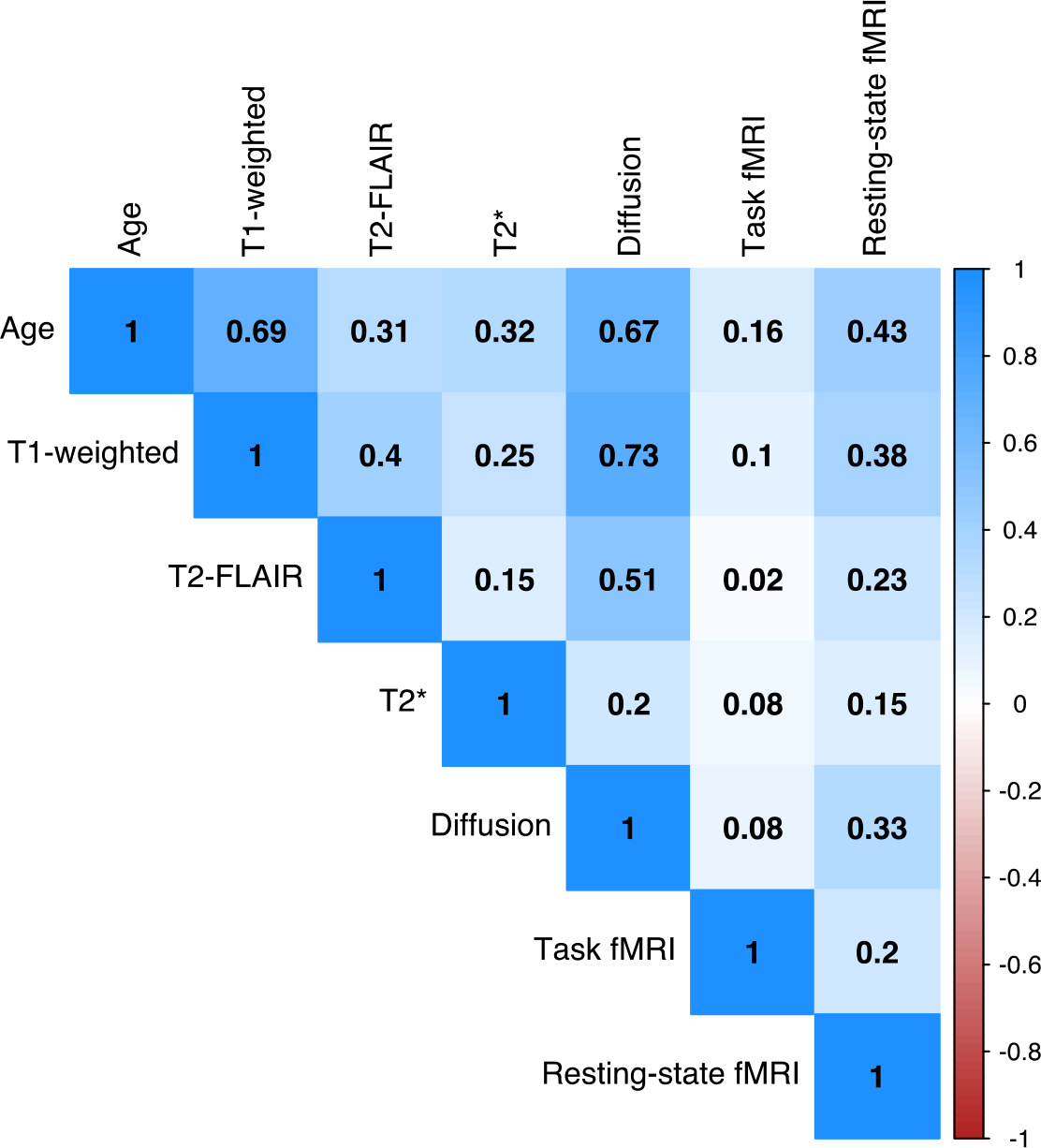
Correlation matrix of age and brain-age predicted by six different modalities. Bivariate correlations between chronological age and brain-age values derived from each of the six neuroimaging modalities, in the validation set (n=520). Values are Pearson’s r for each pairwise correlation. Darker blue colours indicate higher positive correlations.

### 3.4. Brain predicted-age, health, lifestyle and cognitive performance

When applied to the n=14,701 test dataset participants from UK Biobank, performance was similar to that in the validation set: r=0.806, R_2_=0.649, MAE=3.523 years (Figure 3A). A pronounced age bias was apparent (r=−0.624), which was adjusted for using the slope (0.59) and intercept (24.7) of the age on brain-predicted age relationship. After adjustment the age bias was minimal (Figure 3B).

**Figure 3.**
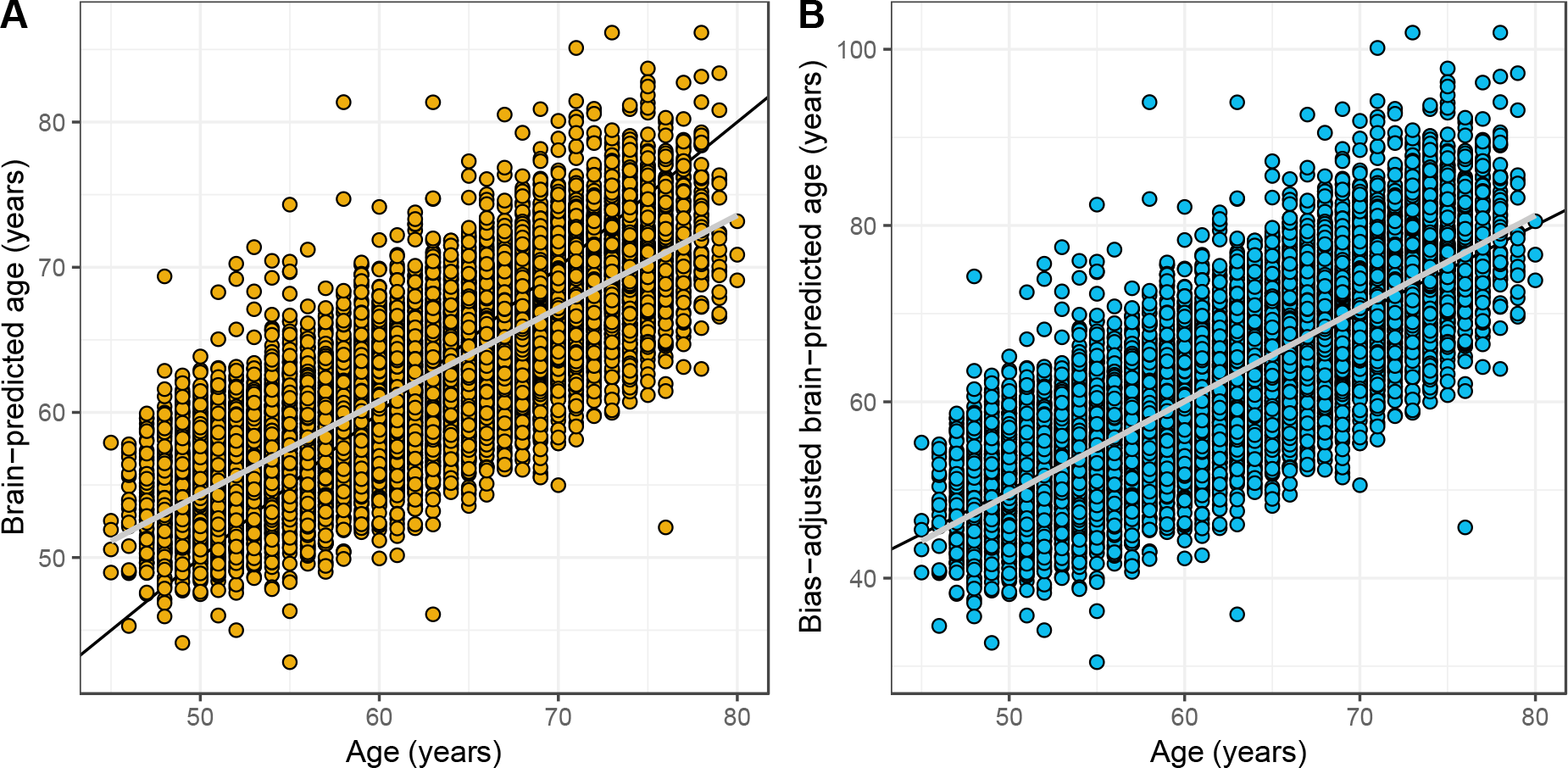
Brain-predicted age by chronological age in the test set, with and without age-bias adjustment. Scatterplots depicting chronological age (x-axis) by brain-predicted age (y-axis) in UK Biobank test set (n=14,701). Black line is the line of identity. Grey line is the regression line of age on brain-predicted age with shaded areas representing the 95% confidence intervals. A) Brain-predicted age values generated from application of previously trained LASSO regression model. The age bias is evident from the slope of the regression line. B) Brain-predicted age values have been adjusted by the slope and intercept of the age-bias line in the training set.

Using the bias-adjusted brain-PAD values, the relationships with the selected outcome measures were assessed (Table 4). Each analysis used a linear regression model, adjusting for age, age_2_, sex, height, volumetric T1-weighted MRI scaling and head motion. Resulting p-values were false-discovery rate corrected (18 tests). After multiple comparison correction, increasing brain-PAD was associated with: higher blood pressure, a diagnosis of diabetes, a history of stroke, past or present smoking, greater frequency of alcohol intake, lower fluid intelligence, longer duration to complete the alphanumeric path trail 2, fewer correct matrix pattern puzzles complete and fewer correct Tower rearranging puzzles. There was no association with anthropometric measures, facial ageing, physical activity levels and some of the cognitive tasks.

**Table 4.**
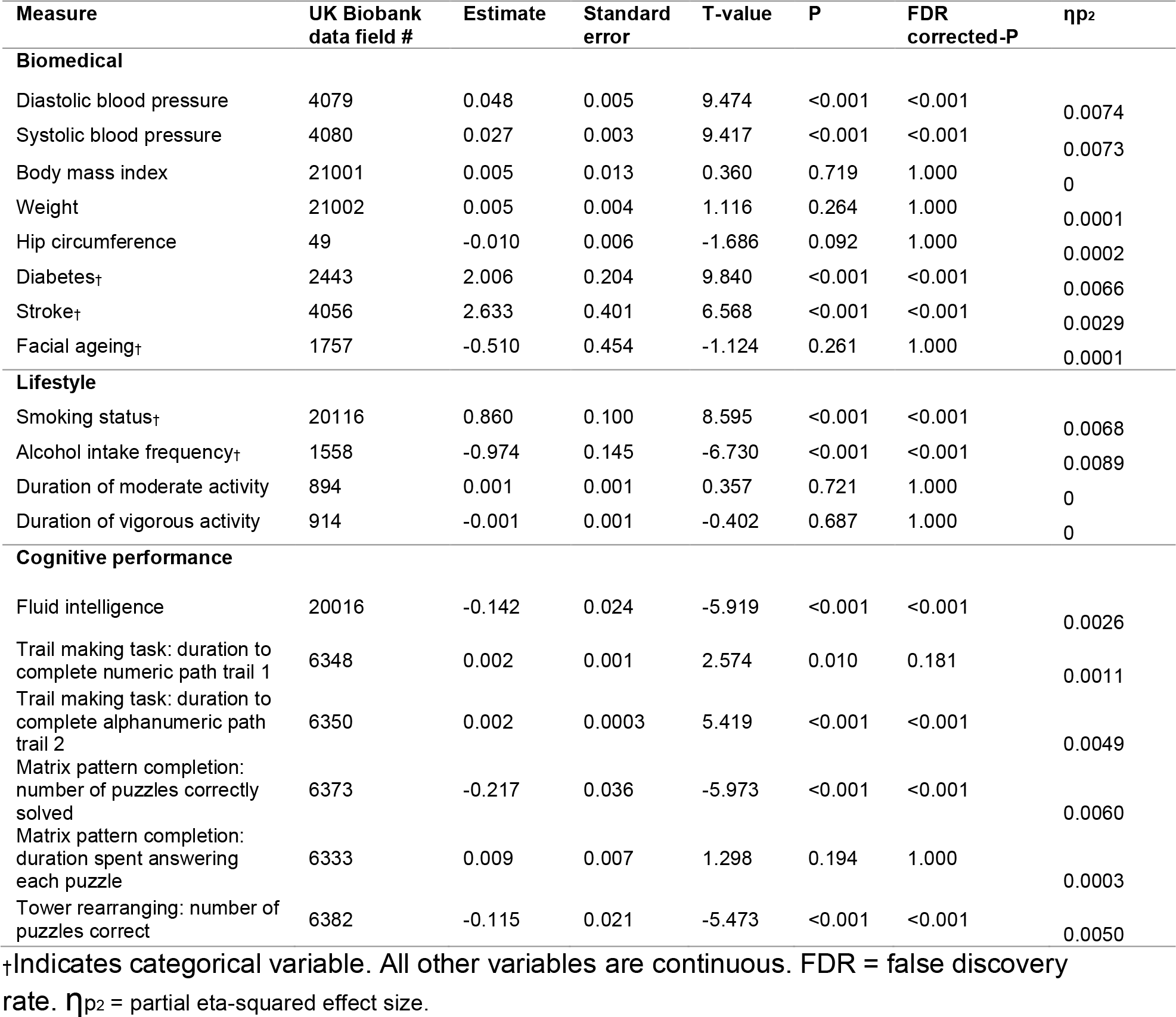
Biomedical, lifestyle and cognitive measures in relation to brain-age.

## 4. Discussion

Using UK Biobank, chronological age can be accurately predicted in healthy people by combining data from six different neuroimaging modalities (T1-weighted MRI, T2-FLAIR, T2*, diffusion-MRI, task fMRI, resting-state fMRI). T1-weighted MRI and diffusion-MRI phenotypes were generally the most informative for age prediction in this combined model. Bootstrapping highlighted 34 variables as informative, and the multi-modality model outperformed prediction models from any modality independently, as hypothesised. This indicates that much of the age-related variation can be captured by these 34 phenotypes alone. Of the independent modality predictors, only T1-weighted MRI and diffusion-MRI achieved reasonable prediction accuracy. Both these modalities performed similarly, contrary to the expectation that T1-weighted MRI would provide the most accurate separate model. When applying the multi-modality model to held-out test data, aberrant brain ageing (quantified by brain-PAD) was associated with higher blood pressure, a history of stroke, diabetes, smoking, frequent alcohol intake and poor cognitive performance.

The majority of extant brain-age studies use T1-weighted MRI alone (Cole, et al., 2019b), though previous multi-modality studies have used two or three modalities (Cherubini, et al., 2016, Groves, et al., 2012, Liem, et al., 2017, Richard, et al., 2018). Thanks to UK Biobank, I was able to combine and compared six different modalities As anticipated, T1-weighted MRI proved important for brain-age prediction here, with normalised grey matter volume being the most informative neuroimaging phenotype. Diffusion-MRI phenotypes were also informative in the LASSO model, although T2* in the putamen, BOLD response during the faces and shapes tasks, and independent components derived from resting-state fMRI were also included. The use of 1000 bootstraps to derive confidence intervals provides assurance of the robustness of the selected features, despite the relatively small contributory effects of each of the 34 informative phenotypes. Interestingly, the predictive performance of the multi-modality model (*r* = 0.79) was lower than some previous reports using multiple modalities (e.g., Cherubini et al., *r* = 0.96), despite the much larger training sample size (n = 2,205 versus n = 140). The current use of derived summary measures (or imaging-derived phenotypes) may negatively impact brain-age prediction performance since these summary measures collapse complex high-dimensional information into single values, potentially removing considerable amounts of age-related variance. In fact, brain-age prediction performance using voxelwise T1-weighted MRI data alone tends to be highly accurately (e.g., r = 0.96; Cole, et al., 2017a). Future work will utilise the raw neuroimaging data from each UK Biobank modality to fuse multi-modality data in a more high-dimensional manner.

One important finding of the current study is that although many of the 1,079 neuroimaging phenotypes significantly correlated with age, with this large sample (n = 2,725) correlations with *r* > 0.05 will be significant at p < 0.05. Hence, many of these statistically significant effects are so small as to be negligible, and only 34 phenotypes were selected as being informative for age-prediction. This means that although multiple patterns of brain structure and function are statistically associated with age, only a few measures are potentially suitably for deriving individualised brain-age predictions with sufficient accuracy for clinical utility. For brain-age, as with other research, it is important to be mindful of the differences between statistical significance, effect size and predictive power in new data.

Another noteworthy issue is the validity of the processing methods for deriving the neuroimaging phenotypes. For example, in the current study subcortical grey matters volume phenotypes were generated in two ways; using FSL FIRST (Patenaude, et al., 2011) and FSL FAST (Zhang, et al., 2001), the latter with a subcortical regional mask applied to the tissue-segmented image. Despite theoretically measuring the same underlying construct, the correlation between FIRST thalamus volume and FAST thalamus volume, for example, is only *r* = 0.44 and *r* = 0.41 for the left and right respectively. This lack of internal consistency highlights the need for researchers using ‘big data’ resources like UK Biobank to carefully consider the nuances of the data processing and to select phenotypes explicitly, as opposed to blindly analysing all available data. This is an important caveat when interpreting the current and other works using these data. Proper validation will come from not only using truly independent datasets but also different processing methods in order to converge towards the same constructs (i.e., brain region segmentation), thus overcoming the assumptions and errors of each separate method.

By using healthy people only for training, a new individual’s deviation from this healthy brain ageing model can be indexed. This key point makes a distinct between models trained on broad inclusive population samples and models trained specifically on healthy people. With the latter, the goal is always to improve model prediction, while with the former, improved model prediction may mean that anyone’s age can be accurately predicted, but the concept of deviation loses value. Previous research has not always screened for healthy people model training, or at least has not reported that this was the case (Smith, et al., 2019, Sun, et al., 2019). This means that potentially the brain-age difference metrics (i.e., brain-age gap, brain-age delta) do not reflect actual deviations from healthy brain ageing and thus may be less sensitive to subsequent relationships with other measures or more prone to false positives. The current study has a large sample with detailed biomedical data, allowing thorough screening of the healthy training dataset. While it is true that even some of these healthy participants may have undetected or prodromal pathologies, this is the case for all case-control research and can only be properly assessed using long-term follow-up. This will also be possible in future thanks to the study design of UK Biobank.

An older-appearing brain was associated with higher diastolic and systolic blood pressure, a history of stroke, a diagnosis of diabetes, smoking, alcohol intake and some facets of cognitive performance. BMI, weight, hip circumference, facial ageing and other aspects of cognitive performance were not associated with brain-PAD. These findings partially concur with previous reports. Franke and colleagues (2014) reported brainAGE (equivalent to brain-PAD) associations with diastolic blood pressure and BMI; here only the former was replicated. Obesity has previously been linked with added brain-ageing (Kolenic, et al., 2018, Ronan, et al., 2016), and BMI has been shown to influence brain structure (Cole, et al., 2013), though no BMI and brain-PAD association was found here. Exercise duration was not associated with brain-PAD, contrary to the findings of Steffener and colleagues (2016). Nevertheless, many different approaches to measuring physical activity are possible, with the current self-report measures not necessarily being the most valid.

Regarding medical history, Franke and colleagues (2013) found that diabetes was associated with an increased brainAGE = 4.6 years, whereas here diabetes increased brain-PAD by 2 years (after covariate adjustment). Stroke was also associated with an increased brain-PAD = 2.6 years; interestingly neither diabetes nor stroke were associated with brain-PAD in the Lothian Birth Cohort 1936 (Cole, et al., 2018). Discrepancies between the current previous work may be due to sampling biases, such as cohort effects or recruitment strategies, though statistical power is certainly in favour of the current study thanks to UK Biobank’s large sample size. Another reason may be the analytic strategy used, particularly the use of covariates. Here, multiple covariates were used to increase sensitivity to brain-PAD relationships, most notably head motion (averaged across the task fMRI session), which accounted for 4.7% of the variance in brain-PAD. While several biomedical, lifestyle and cognitive factors relate to brain-PAD, effect sizes are generally small. For instance, the effects of diabetes, stroke, current smoking (+2.1 years brain-PAD) and daily alcohol intake (+0.97 years) are substantially smaller than previously reported effects of Alzheimer’s (+10 years) or multiple sclerosis (+11 years) on apparent brain-age (Cole, et al., 2019a, Franke, et al., 2010). While it makes intuitive sense that such factors are associated with poorer ageing-related brain health, the magnitude of the effects should not be overstated. Future work will investigate gene-environment interactions to better understand individual differences in the impact of medical history and lifestyle on brain health during ageing.

Fluid intelligence has previously been linked to brain-PAD (Cole, et al., 2018), as has performance on the trail-making task (Cole, et al., 2015, Cole, et al., 2017b), and other studies report moderate significant relationships between brain-ageing and cognitive performance (Liem, et al., 2017, Richard, et al., 2018). Here I was able to replicate that relationship, though more detailed analysis of how brain-ageing relates to cognitive ageing will require more comprehensive cognitive testing. The presence of data on facial ageing presented an interesting opportunity to test the relationship between brain-PAD and self-report of whether participants felt people said they looked “younger than you are”, “older than you are” or “about the same age”. There was no relationship detected, although interesting related work has reported a relationship between subjective “age” and brain-age (Kwak, et al., 2018). In the current study, only 1% of respondents reported that people think they look older, potentially questioning the validity of this subjective measure.

The current study benefits from the extremely large sample size and in-depth biomedical data of UK Biobank. This enabled detailed screening of healthy individuals to include in the training dataset, to minimise the impact of disease-related effects on the brain that may potentially confound models of healthy brain ageing. Despite 84% of the neuroimaging sample from UK Biobank used herein having at least one ICD-10 diagnosis, a self-reported longstanding illness, poor self-rated health status or medical history of stroke or diabetes, the remaining 16% still left greater than 2,500 participants for training and validation. Another benefit here is the use of LASSO regression with bootstrapping, combining the strength of LASSO for identifying important features for prediction (by shrinking uninformative variables to zero) with the robustness of bootstrapping, which overcomes the limitation of using LASSO with highly correlated predictor variables. A weakness of the study is the use of summary-level neuroimaging phenotypes. As noted, model accuracy is substantially below that commonly achieved with voxelwise data. Another potential limitation is the current limited use of biomedical, lifestyle and cognitive variables. Potentially many more associations with biological, psychological and behavioural parameters could have been assessed, however adding further tests adds to the multiple-testing burden and decreases sensitivity to real effects. Hence the decision was taken to test only variables with previous research evidence and strong face validity. Another weakness is that the current study is cross-sectional, hence the long-term consequences of having an older-appearing brain cannot be tested and no causal inference made. The planned long-term medical follow-up of UK Biobank participants makes this an obvious avenue for future research.

## 5. Conclusions

In summary, brain-age can be predicted by combining six different MRI modalities, with the strongest predictors being T1-weighted and diffusion-MRI phenotypes. Deviations from healthy brain ageing are related to medical history, smoking and alcohol intake. Poorer cognitive performance was also related to having an older-appearing brain, suggesting that biomedical and lifestyle factors can negatively impact both brain ageing and cognitive ageing. The brain-age paradigm presents a potential screening tool to detect the negative impact of medical and lifestyle factors on brain health in asymptomatic people, and the planned longitudinal nature of UK Biobank offers the opportunity to validate this potential.

## Acknowledgements

The author would like to thank the UK Biobank participants for giving up their time for this unique project.

