## Supplementary material for "Multi-modality neuroimaging brain-age in UK Biobank: relationship to biomedical, lifestyle and cognitive factors": Table A1

**Appendix**

**Table A1. UK Biobank neuroimaging phenotypes**

| **Neuroimaging phenotype** | **UK Biobank data field #** |
| --- | --- |
| Volumetric scaling from T1 head image to standard space | 25000 |
| Volume of peripheral cortical grey matter normalised for head size | 25001 |
| Volume of peripheral cortical grey matter | 25002 |
| Volume of ventricular cerebrospinal fluid normalised for head size | 25003 |
| Volume of ventricular cerebrospinal fluid | 25004 |
| Volume of grey matter normalised for head size | 25005 |
| Volume of grey matter | 25006 |
| Volume of white matter normalised for head size | 25007 |
| Volume of white matter | 25008 |
| Volume of brain grey + white matter normalised for head size | 25009 |
| Volume of brain grey + white matter | 25010 |
| Volume of thalamus left | 25011 |
| Volume of thalamus right | 25012 |
| Volume of caudate left | 25013 |
| Volume of caudate right | 25014 |
| Volume of putamen left | 25015 |
| Volume of putamen right | 25016 |
| Volume of pallidum left | 25017 |
| Volume of pallidum right | 25018 |
| Volume of hippocampus left | 25019 |
| Volume of hippocampus right | 25020 |
| Volume of amygdala left | 25021 |
| Volume of amygdala right | 25022 |
| Volume of accumbens left | 25023 |
| Volume of accumbens right | 25024 |
| Volume of brain stem 4th ventricle | 25025 |
| Median T2* in thalamus left | 25026 |
| Median T2* in thalamus right | 25027 |
| Median T2* in caudate left | 25028 |
| Median T2* in caudate right | 25029 |
| Median T2* in putamen left | 25030 |
| Median T2* in putamen right | 25031 |
| Median T2* in pallidum left | 25032 |
| Median T2* in pallidum right | 25033 |
| Median T2* in hippocampus left | 25034 |
| Median T2* in hippocampus right | 25035 |
| Median T2* in amygdala left | 25036 |
| Median T2* in amygdala right | 25037 |
| Median T2* in accumbens left | 25038 |
| Median T2* in accumbens right | 25039 |
| Median BOLD effect in group-defined mask for shapes activation | 25040 |
| Median Z-statistic in group-defined mask for shapes activation | 25042 |
| Median BOLD effect in group-defined mask for faces activation | 25044 |
| Median Z-statistic in group-defined mask for faces activation | 25046 |
| Median BOLD effect in group-defined mask for faces-shapes contrast | 25048 |
| Median BOLD effect in group-defined amygdala activation mask for faces-shapes contrast | 25052 |
| Median Z-statistic in group-defined amygdala activation mask for faces-shapes contrast | 25054 |
| 90th percentile of BOLD effect in group-defined mask for shapes activation | 25761 |
| 90th percentile of Z-statistic in group-defined mask for shapes activation | 25762 |
| 90th percentile of BOLD effect in group-defined mask for faces activation | 25763 |
| 90th percentile of Z-statistic in group-defined mask for faces activation | 25764 |
| 90th percentile of BOLD effect in group-defined mask for faces-shapes contrast | 25765 |
| 90th percentile of BOLD effect in group-defined amygdala activation mask for faces-shapes contrast | 25767 |
| 90th percentile of Z-statistic in group-defined amygdala activation mask for faces-shapes contrast | 25768 |
| Total volume of white matter hyperintensities from T1-weighted and T2-FLAIR images | 25781 |
| Volume of grey matter in frontal pole left | 25782 |
| Volume of grey matter in frontal pole right | 25783 |
| Volume of grey matter in insular cortex left | 25784 |
| Volume of grey matter in insular cortex right | 25785 |
| Volume of grey matter in superior frontal gyrus left | 25786 |
| Volume of grey matter in superior frontal gyrus right | 25787 |
| Volume of grey matter in middle frontal gyrus left | 25788 |
| Volume of grey matter in middle frontal gyrus right | 25789 |
| Volume of grey matter in inferior frontal gyrus pars triangularis left | 25790 |
| Volume of grey matter in inferior frontal gyrus pars triangularis right | 25791 |
| Volume of grey matter in inferior frontal gyrus pars opercularis left | 25792 |
| Volume of grey matter in inferior frontal gyrus pars opercularis right | 25793 |
| Volume of grey matter in precentral gyrus left | 25794 |
| Volume of grey matter in precentral gyrus right | 25795 |
| Volume of grey matter in temporal pole left | 25796 |
| Volume of grey matter in temporal pole right | 25797 |
| Volume of grey matter in superior temporal gyrus anterior division left | 25798 |
| Volume of grey matter in superior temporal gyrus anterior division right | 25799 |
| Volume of grey matter in superior temporal gyrus posterior division left | 25800 |
| Volume of grey matter in superior temporal gyrus posterior division right | 25801 |
| Volume of grey matter in middle temporal gyrus anterior division left | 25802 |
| Volume of grey matter in middle temporal gyrus anterior division right | 25803 |
| Volume of grey matter in middle temporal gyrus posterior division left | 25804 |
| Volume of grey matter in middle temporal gyrus posterior division right | 25805 |
| Volume of grey matter in middle temporal gyrus temporooccipital part left | 25806 |
| Volume of grey matter in middle temporal gyrus temporooccipital part right | 25807 |
| Volume of grey matter in inferior temporal gyrus anterior division left | 25808 |
| Volume of grey matter in inferior temporal gyrus anterior division right | 25809 |
| Volume of grey matter in inferior temporal gyrus posterior division left | 25810 |
| Volume of grey matter in inferior temporal gyrus posterior division right | 25811 |
| Volume of grey matter in inferior temporal gyrus temporooccipital part left | 25812 |
| Volume of grey matter in inferior temporal gyrus temporooccipital part right | 25813 |
| Volume of grey matter in postcentral gyrus left | 25814 |
| Volume of grey matter in postcentral gyrus right | 25815 |
| Volume of grey matter in superior parietal lobule left | 25816 |
| Volume of grey matter in superior parietal lobule right | 25817 |
| Volume of grey matter in supramarginal gyrus anterior division left | 25818 |
| Volume of grey matter in supramarginal gyrus anterior division right | 25819 |
| Volume of grey matter in supramarginal gyrus posterior division left | 25820 |
| Volume of grey matter in supramarginal gyrus posterior division right | 25821 |
| Volume of grey matter in angular gyrus left | 25822 |
| Volume of grey matter in angular gyrus right | 25823 |
| Volume of grey matter in lateral occipital cortex superior division left | 25824 |
| Volume of grey matter in lateral occipital cortex superior division right | 25825 |
| Volume of grey matter in lateral occipital cortex inferior division left | 25826 |
| Volume of grey matter in lateral occipital cortex inferior division right | 25827 |
| Volume of grey matter in intracalcarine cortex left | 25828 |
| Volume of grey matter in intracalcarine cortex right | 25829 |
| Volume of grey matter in frontal medial cortex left | 25830 |
| Volume of grey matter in frontal medial cortex right | 25831 |
| Volume of grey matter in juxtapositional lobule cortex formerly supplementary motor cortex left | 25832 |
| Volume of grey matter in juxtapositional lobule cortex formerly supplementary motor cortex right | 25833 |
| Volume of grey matter in subcallosal cortex left | 25834 |
| Volume of grey matter in subcallosal cortex right | 25835 |
| Volume of grey matter in paracingulate gyrus left | 25836 |
| Volume of grey matter in paracingulate gyrus right | 25837 |
| Volume of grey matter in cingulate gyrus anterior division left | 25838 |
| Volume of grey matter in cingulate gyrus anterior division right | 25839 |
| Volume of grey matter in cingulate gyrus posterior division left | 25840 |
| Volume of grey matter in cingulate gyrus posterior division right | 25841 |
| Volume of grey matter in precuneous cortex left | 25842 |
| Volume of grey matter in precuneous cortex right | 25843 |
| Volume of grey matter in cuneal cortex left | 25844 |
| Volume of grey matter in cuneal cortex right | 25845 |
| Volume of grey matter in frontal orbital cortex left | 25846 |
| Volume of grey matter in frontal orbital cortex right | 25847 |
| Volume of grey matter in parahippocampal gyrus anterior division left | 25848 |
| Volume of grey matter in parahippocampal gyrus anterior division right | 25849 |
| Volume of grey matter in parahippocampal gyrus posterior division left | 25850 |
| Volume of grey matter in parahippocampal gyrus posterior division right | 25851 |
| Volume of grey matter in lingual gyrus left | 25852 |
| Volume of grey matter in lingual gyrus right | 25853 |
| Volume of grey matter in temporal fusiform cortex anterior division left | 25854 |
| Volume of grey matter in temporal fusiform cortex anterior division right | 25855 |
| Volume of grey matter in temporal fusiform cortex posterior division left | 25856 |
| Volume of grey matter in temporal fusiform cortex posterior division right | 25857 |
| Volume of grey matter in temporal occipital fusiform cortex left | 25858 |
| Volume of grey matter in temporal occipital fusiform cortex right | 25859 |
| Volume of grey matter in occipital fusiform gyrus left | 25860 |
| Volume of grey matter in occipital fusiform gyrus right | 25861 |
| Volume of grey matter in frontal operculum cortex left | 25862 |
| Volume of grey matter in frontal operculum cortex right | 25863 |
| Volume of grey matter in central opercular cortex left | 25864 |
| Volume of grey matter in central opercular cortex right | 25865 |
| Volume of grey matter in parietal operculum cortex left | 25866 |
| Volume of grey matter in parietal operculum cortex right | 25867 |
| Volume of grey matter in planum polare left | 25868 |
| Volume of grey matter in planum polare right | 25869 |
| Volume of grey matter in heschls gyrus includes h1 and h2 left | 25870 |
| Volume of grey matter in heschls gyrus includes h1 and h2 right | 25871 |
| Volume of grey matter in planum temporale left | 25872 |
| Volume of grey matter in planum temporale right | 25873 |
| Volume of grey matter in supracalcarine cortex left | 25874 |
| Volume of grey matter in supracalcarine cortex right | 25875 |
| Volume of grey matter in occipital pole left | 25876 |
| Volume of grey matter in occipital pole right | 25877 |
| Volume of grey matter in thalamus left | 25878 |
| Volume of grey matter in thalamus right | 25879 |
| Volume of grey matter in caudate left | 25880 |
| Volume of grey matter in caudate right | 25881 |
| Volume of grey matter in putamen left | 25882 |
| Volume of grey matter in putamen right | 25883 |
| Volume of grey matter in pallidum left | 25884 |
| Volume of grey matter in pallidum right | 25885 |
| Volume of grey matter in hippocampus left | 25886 |
| Volume of grey matter in hippocampus right | 25887 |
| Volume of grey matter in amygdala left | 25888 |
| Volume of grey matter in amygdala right | 25889 |
| Volume of grey matter in ventral striatum left | 25890 |
| Volume of grey matter in ventral striatum right | 25891 |
| Volume of grey matter in brainstem | 25892 |
| Volume of grey matter in iiv cerebellum left | 25893 |
| Volume of grey matter in iiv cerebellum right | 25894 |
| Volume of grey matter in v cerebellum left | 25895 |
| Volume of grey matter in v cerebellum right | 25896 |
| Volume of grey matter in vi cerebellum left | 25897 |
| Volume of grey matter in vermis vi cerebellum | 25898 |
| Volume of grey matter in vi cerebellum right | 25899 |
| Volume of grey matter in crus i cerebellum left | 25900 |
| Volume of grey matter in vermis crus i cerebellum | 25901 |
| Volume of grey matter in crus i cerebellum right | 25902 |
| Volume of grey matter in crus ii cerebellum left | 25903 |
| Volume of grey matter in vermis crus ii cerebellum | 25904 |
| Volume of grey matter in crus ii cerebellum right | 25905 |
| Volume of grey matter in viib cerebellum left | 25906 |
| Volume of grey matter in vermis viib cerebellum | 25907 |
| Volume of grey matter in viib cerebellum right | 25908 |
| Volume of grey matter in viiia cerebellum left | 25909 |
| Volume of grey matter in vermis viiia cerebellum | 25910 |
| Volume of grey matter in viiia cerebellum right | 25911 |
| Volume of grey matter in viiib cerebellum left | 25912 |
| Volume of grey matter in vermis viiib cerebellum | 25913 |
| Volume of grey matter in viiib cerebellum right | 25914 |
| Volume of grey matter in ix cerebellum left | 25915 |
| Volume of grey matter in vermis ix cerebellum | 25916 |
| Volume of grey matter in ix cerebellum right | 25917 |
| Volume of grey matter in x cerebellum left | 25918 |
| Volume of grey matter in vermis x cerebellum | 25919 |
| Volume of grey matter in x cerebellum right | 25920 |
| Mean FA in middle cerebellar peduncle on FA skeleton | 25056 |
| Mean FA in pontine crossing tract on FA skeleton | 25057 |
| Mean FA in genu of corpus callosum on FA skeleton | 25058 |
| Mean FA in body of corpus callosum on FA skeleton | 25059 |
| Mean FA in splenium of corpus callosum on FA skeleton | 25060 |
| Mean FA in fornix on FA skeleton | 25061 |
| Mean FA in corticospinal tract on FA skeleton right | 25062 |
| Mean FA in corticospinal tract on FA skeleton left | 25063 |
| Mean FA in medial lemniscus on FA skeleton right | 25064 |
| Mean FA in medial lemniscus on FA skeleton left | 25065 |
| Mean FA in inferior cerebellar peduncle on FA skeleton right | 25066 |
| Mean FA in inferior cerebellar peduncle on FA skeleton left | 25067 |
| Mean FA in superior cerebellar peduncle on FA skeleton right | 25068 |
| Mean FA in superior cerebellar peduncle on FA skeleton left | 25069 |
| Mean FA in cerebral peduncle on FA skeleton right | 25070 |
| Mean FA in cerebral peduncle on FA skeleton left | 25071 |
| Mean FA in anterior limb of internal capsule on FA skeleton right | 25072 |
| Mean FA in anterior limb of internal capsule on FA skeleton left | 25073 |
| Mean FA in posterior limb of internal capsule on FA skeleton right | 25074 |
| Mean FA in posterior limb of internal capsule on FA skeleton left | 25075 |
| Mean FA in retrolenticular part of internal capsule on FA skeleton right | 25076 |
| Mean FA in retrolenticular part of internal capsule on FA skeleton left | 25077 |
| Mean FA in anterior corona radiata on FA skeleton right | 25078 |
| Mean FA in anterior corona radiata on FA skeleton left | 25079 |
| Mean FA in superior corona radiata on FA skeleton right | 25080 |
| Mean FA in superior corona radiata on FA skeleton left | 25081 |
| Mean FA in posterior corona radiata on FA skeleton right | 25082 |
| Mean FA in posterior corona radiata on FA skeleton left | 25083 |
| Mean FA in posterior thalamic radiation on FA skeleton right | 25084 |
| Mean FA in posterior thalamic radiation on FA skeleton left | 25085 |
| Mean FA in sagittal stratum on FA skeleton right | 25086 |
| Mean FA in sagittal stratum on FA skeleton left | 25087 |
| Mean FA in external capsule on FA skeleton right | 25088 |
| Mean FA in external capsule on FA skeleton left | 25089 |
| Mean FA in cingulum cingulate gyrus on FA skeleton right | 25090 |
| Mean FA in cingulum cingulate gyrus on FA skeleton left | 25091 |
| Mean FA in cingulum hippocampus on FA skeleton right | 25092 |
| Mean FA in cingulum hippocampus on FA skeleton left | 25093 |
| Mean FA in fornix cres/stria terminalis on FA skeleton right | 25094 |
| Mean FA in fornix cres/stria terminalis on FA skeleton left | 25095 |
| Mean FA in superior longitudinal fasciculus on FA skeleton right | 25096 |
| Mean FA in superior longitudinal fasciculus on FA skeleton left | 25097 |
| Mean FA in superior frontooccipital fasciculus on FA skeleton right | 25098 |
| Mean FA in superior frontooccipital fasciculus on FA skeleton left | 25099 |
| Mean FA in uncinate fasciculus on FA skeleton right | 25100 |
| Mean FA in uncinate fasciculus on FA skeleton left | 25101 |
| Mean FA in tapetum on FA skeleton right | 25102 |
| Mean FA in tapetum on FA skeleton left | 25103 |
| Mean MD in middle cerebellar peduncle on FA skeleton | 25104 |
| Mean MD in pontine crossing tract on FA skeleton | 25105 |
| Mean MD in genu of corpus callosum on FA skeleton | 25106 |
| Mean MD in body of corpus callosum on FA skeleton | 25107 |
| Mean MD in splenium of corpus callosum on FA skeleton | 25108 |
| Mean MD in fornix on FA skeleton | 25109 |
| Mean MD in corticospinal tract on FA skeleton right | 25110 |
| Mean MD in corticospinal tract on FA skeleton left | 25111 |
| Mean MD in medial lemniscus on FA skeleton right | 25112 |
| Mean MD in medial lemniscus on FA skeleton left | 25113 |
| Mean MD in inferior cerebellar peduncle on FA skeleton right | 25114 |
| Mean MD in inferior cerebellar peduncle on FA skeleton left | 25115 |
| Mean MD in superior cerebellar peduncle on FA skeleton right | 25116 |
| Mean MD in superior cerebellar peduncle on FA skeleton left | 25117 |
| Mean MD in cerebral peduncle on FA skeleton right | 25118 |
| Mean MD in cerebral peduncle on FA skeleton left | 25119 |
| Mean MD in anterior limb of internal capsule on FA skeleton right | 25120 |
| Mean MD in anterior limb of internal capsule on FA skeleton left | 25121 |
| Mean MD in posterior limb of internal capsule on FA skeleton right | 25122 |
| Mean MD in posterior limb of internal capsule on FA skeleton left | 25123 |
| Mean MD in retrolenticular part of internal capsule on FA skeleton right | 25124 |
| Mean MD in retrolenticular part of internal capsule on FA skeleton left | 25125 |
| Mean MD in anterior corona radiata on FA skeleton right | 25126 |
| Mean MD in anterior corona radiata on FA skeleton left | 25127 |
| Mean MD in superior corona radiata on FA skeleton right | 25128 |
| Mean MD in superior corona radiata on FA skeleton left | 25129 |
| Mean MD in posterior corona radiata on FA skeleton right | 25130 |
| Mean MD in posterior corona radiata on FA skeleton left | 25131 |
| Mean MD in posterior thalamic radiation on FA skeleton right | 25132 |
| Mean MD in posterior thalamic radiation on FA skeleton left | 25133 |
| Mean MD in sagittal stratum on FA skeleton right | 25134 |
| Mean MD in sagittal stratum on FA skeleton left | 25135 |
| Mean MD in external capsule on FA skeleton right | 25136 |
| Mean MD in external capsule on FA skeleton left | 25137 |
| Mean MD in cingulum cingulate gyrus on FA skeleton right | 25138 |
| Mean MD in cingulum cingulate gyrus on FA skeleton left | 25139 |
| Mean MD in cingulum hippocampus on FA skeleton right | 25140 |
| Mean MD in cingulum hippocampus on FA skeleton left | 25141 |
| Mean MD in fornix cres/stria terminalis on FA skeleton right | 25142 |
| Mean MD in fornix cres/stria terminalis on FA skeleton left | 25143 |
| Mean MD in superior longitudinal fasciculus on FA skeleton right | 25144 |
| Mean MD in superior longitudinal fasciculus on FA skeleton left | 25145 |
| Mean MD in superior frontooccipital fasciculus on FA skeleton right | 25146 |
| Mean MD in superior frontooccipital fasciculus on FA skeleton left | 25147 |
| Mean MD in uncinate fasciculus on FA skeleton right | 25148 |
| Mean MD in uncinate fasciculus on FA skeleton left | 25149 |
| Mean MD in tapetum on FA skeleton right | 25150 |
| Mean MD in tapetum on FA skeleton left | 25151 |
| Mean MO in middle cerebellar peduncle on FA skeleton | 25152 |
| Mean MO in pontine crossing tract on FA skeleton | 25153 |
| Mean MO in genu of corpus callosum on FA skeleton | 25154 |
| Mean MO in body of corpus callosum on FA skeleton | 25155 |
| Mean MO in splenium of corpus callosum on FA skeleton | 25156 |
| Mean MO in fornix on FA skeleton | 25157 |
| Mean MO in corticospinal tract on FA skeleton right | 25158 |
| Mean MO in corticospinal tract on FA skeleton left | 25159 |
| Mean MO in medial lemniscus on FA skeleton right | 25160 |
| Mean MO in medial lemniscus on FA skeleton left | 25161 |
| Mean MO in inferior cerebellar peduncle on FA skeleton right | 25162 |
| Mean MO in inferior cerebellar peduncle on FA skeleton left | 25163 |
| Mean MO in superior cerebellar peduncle on FA skeleton right | 25164 |
| Mean MO in superior cerebellar peduncle on FA skeleton left | 25165 |
| Mean MO in cerebral peduncle on FA skeleton right | 25166 |
| Mean MO in cerebral peduncle on FA skeleton left | 25167 |
| Mean MO in anterior limb of internal capsule on FA skeleton right | 25168 |
| Mean MO in anterior limb of internal capsule on FA skeleton left | 25169 |
| Mean MO in posterior limb of internal capsule on FA skeleton right | 25170 |
| Mean MO in posterior limb of internal capsule on FA skeleton left | 25171 |
| Mean MO in retrolenticular part of internal capsule on FA skeleton right | 25172 |
| Mean MO in retrolenticular part of internal capsule on FA skeleton left | 25173 |
| Mean MO in anterior corona radiata on FA skeleton right | 25174 |
| Mean MO in anterior corona radiata on FA skeleton left | 25175 |
| Mean MO in superior corona radiata on FA skeleton right | 25176 |
| Mean MO in superior corona radiata on FA skeleton left | 25177 |
| Mean MO in posterior corona radiata on FA skeleton right | 25178 |
| Mean MO in posterior corona radiata on FA skeleton left | 25179 |
| Mean MO in posterior thalamic radiation on FA skeleton right | 25180 |
| Mean MO in posterior thalamic radiation on FA skeleton left | 25181 |
| Mean MO in sagittal stratum on FA skeleton right | 25182 |
| Mean MO in sagittal stratum on FA skeleton left | 25183 |
| Mean MO in external capsule on FA skeleton right | 25184 |
| Mean MO in external capsule on FA skeleton left | 25185 |
| Mean MO in cingulum cingulate gyrus on FA skeleton right | 25186 |
| Mean MO in cingulum cingulate gyrus on FA skeleton left | 25187 |
| Mean MO in cingulum hippocampus on FA skeleton right | 25188 |
| Mean MO in cingulum hippocampus on FA skeleton left | 25189 |
| Mean MO in fornix cres/stria terminalis on FA skeleton right | 25190 |
| Mean MO in fornix cres/stria terminalis on FA skeleton left | 25191 |
| Mean MO in superior longitudinal fasciculus on FA skeleton right | 25192 |
| Mean MO in superior longitudinal fasciculus on FA skeleton left | 25193 |
| Mean MO in superior frontooccipital fasciculus on FA skeleton right | 25194 |
| Mean MO in superior frontooccipital fasciculus on FA skeleton left | 25195 |
| Mean MO in uncinate fasciculus on FA skeleton right | 25196 |
| Mean MO in uncinate fasciculus on FA skeleton left | 25197 |
| Mean MO in tapetum on FA skeleton right | 25198 |
| Mean MO in tapetum on FA skeleton left | 25199 |
| Mean L1 in middle cerebellar peduncle on FA skeleton | 25200 |
| Mean L1 in pontine crossing tract on FA skeleton | 25201 |
| Mean L1 in genu of corpus callosum on FA skeleton | 25202 |
| Mean L1 in body of corpus callosum on FA skeleton | 25203 |
| Mean L1 in splenium of corpus callosum on FA skeleton | 25204 |
| Mean L1 in fornix on FA skeleton | 25205 |
| Mean L1 in corticospinal tract on FA skeleton right | 25206 |
| Mean L1 in corticospinal tract on FA skeleton left | 25207 |
| Mean L1 in medial lemniscus on FA skeleton right | 25208 |
| Mean L1 in medial lemniscus on FA skeleton left | 25209 |
| Mean L1 in inferior cerebellar peduncle on FA skeleton right | 25210 |
| Mean L1 in inferior cerebellar peduncle on FA skeleton left | 25211 |
| Mean L1 in superior cerebellar peduncle on FA skeleton right | 25212 |
| Mean L1 in superior cerebellar peduncle on FA skeleton left | 25213 |
| Mean L1 in cerebral peduncle on FA skeleton right | 25214 |
| Mean L1 in cerebral peduncle on FA skeleton left | 25215 |
| Mean L1 in anterior limb of internal capsule on FA skeleton right | 25216 |
| Mean L1 in anterior limb of internal capsule on FA skeleton left | 25217 |
| Mean L1 in posterior limb of internal capsule on FA skeleton right | 25218 |
| Mean L1 in posterior limb of internal capsule on FA skeleton left | 25219 |
| Mean L1 in retrolenticular part of internal capsule on FA skeleton right | 25220 |
| Mean L1 in retrolenticular part of internal capsule on FA skeleton left | 25221 |
| Mean L1 in anterior corona radiata on FA skeleton right | 25222 |
| Mean L1 in anterior corona radiata on FA skeleton left | 25223 |
| Mean L1 in superior corona radiata on FA skeleton right | 25224 |
| Mean L1 in superior corona radiata on FA skeleton left | 25225 |
| Mean L1 in posterior corona radiata on FA skeleton right | 25226 |
| Mean L1 in posterior corona radiata on FA skeleton left | 25227 |
| Mean L1 in posterior thalamic radiation on FA skeleton right | 25228 |
| Mean L1 in posterior thalamic radiation on FA skeleton left | 25229 |
| Mean L1 in sagittal stratum on FA skeleton right | 25230 |
| Mean L1 in sagittal stratum on FA skeleton left | 25231 |
| Mean L1 in external capsule on FA skeleton right | 25232 |
| Mean L1 in external capsule on FA skeleton left | 25233 |
| Mean L1 in cingulum cingulate gyrus on FA skeleton right | 25234 |
| Mean L1 in cingulum cingulate gyrus on FA skeleton left | 25235 |
| Mean L1 in cingulum hippocampus on FA skeleton right | 25236 |
| Mean L1 in cingulum hippocampus on FA skeleton left | 25237 |
| Mean L1 in fornix cres/stria terminalis on FA skeleton right | 25238 |
| Mean L1 in fornix cres/stria terminalis on FA skeleton left | 25239 |
| Mean L1 in superior longitudinal fasciculus on FA skeleton right | 25240 |
| Mean L1 in superior longitudinal fasciculus on FA skeleton left | 25241 |
| Mean L1 in superior frontooccipital fasciculus on FA skeleton right | 25242 |
| Mean L1 in superior frontooccipital fasciculus on FA skeleton left | 25243 |
| Mean L1 in uncinate fasciculus on FA skeleton right | 25244 |
| Mean L1 in uncinate fasciculus on FA skeleton left | 25245 |
| Mean L1 in tapetum on FA skeleton right | 25246 |
| Mean L1 in tapetum on FA skeleton left | 25247 |
| Mean L2 in middle cerebellar peduncle on FA skeleton | 25248 |
| Mean L2 in pontine crossing tract on FA skeleton | 25249 |
| Mean L2 in genu of corpus callosum on FA skeleton | 25250 |
| Mean L2 in body of corpus callosum on FA skeleton | 25251 |
| Mean L2 in splenium of corpus callosum on FA skeleton | 25252 |
| Mean L2 in fornix on FA skeleton | 25253 |
| Mean L2 in corticospinal tract on FA skeleton right | 25254 |
| Mean L2 in corticospinal tract on FA skeleton left | 25255 |
| Mean L2 in medial lemniscus on FA skeleton right | 25256 |
| Mean L2 in medial lemniscus on FA skeleton left | 25257 |
| Mean L2 in inferior cerebellar peduncle on FA skeleton right | 25258 |
| Mean L2 in inferior cerebellar peduncle on FA skeleton left | 25259 |
| Mean L2 in superior cerebellar peduncle on FA skeleton right | 25260 |
| Mean L2 in superior cerebellar peduncle on FA skeleton left | 25261 |
| Mean L2 in cerebral peduncle on FA skeleton right | 25262 |
| Mean L2 in cerebral peduncle on FA skeleton left | 25263 |
| Mean L2 in anterior limb of internal capsule on FA skeleton right | 25264 |
| Mean L2 in anterior limb of internal capsule on FA skeleton left | 25265 |
| Mean L2 in posterior limb of internal capsule on FA skeleton right | 25266 |
| Mean L2 in posterior limb of internal capsule on FA skeleton left | 25267 |
| Mean L2 in retrolenticular part of internal capsule on FA skeleton right | 25268 |
| Mean L2 in retrolenticular part of internal capsule on FA skeleton left | 25269 |
| Mean L2 in anterior corona radiata on FA skeleton right | 25270 |
| Mean L2 in anterior corona radiata on FA skeleton left | 25271 |
| Mean L2 in superior corona radiata on FA skeleton right | 25272 |
| Mean L2 in superior corona radiata on FA skeleton left | 25273 |
| Mean L2 in posterior corona radiata on FA skeleton right | 25274 |
| Mean L2 in posterior corona radiata on FA skeleton left | 25275 |
| Mean L2 in posterior thalamic radiation on FA skeleton right | 25276 |
| Mean L2 in posterior thalamic radiation on FA skeleton left | 25277 |
| Mean L2 in sagittal stratum on FA skeleton right | 25278 |
| Mean L2 in sagittal stratum on FA skeleton left | 25279 |
| Mean L2 in external capsule on FA skeleton right | 25280 |
| Mean L2 in external capsule on FA skeleton left | 25281 |
| Mean L2 in cingulum cingulate gyrus on FA skeleton right | 25282 |
| Mean L2 in cingulum cingulate gyrus on FA skeleton left | 25283 |
| Mean L2 in cingulum hippocampus on FA skeleton right | 25284 |
| Mean L2 in cingulum hippocampus on FA skeleton left | 25285 |
| Mean L2 in fornix cres/stria terminalis on FA skeleton right | 25286 |
| Mean L2 in fornix cres/stria terminalis on FA skeleton left | 25287 |
| Mean L2 in superior longitudinal fasciculus on FA skeleton right | 25288 |
| Mean L2 in superior longitudinal fasciculus on FA skeleton left | 25289 |
| Mean L2 in superior frontooccipital fasciculus on FA skeleton right | 25290 |
| Mean L2 in superior frontooccipital fasciculus on FA skeleton left | 25291 |
| Mean L2 in uncinate fasciculus on FA skeleton right | 25292 |
| Mean L2 in uncinate fasciculus on FA skeleton left | 25293 |
| Mean L2 in tapetum on FA skeleton right | 25294 |
| Mean L2 in tapetum on FA skeleton left | 25295 |
| Mean L3 in middle cerebellar peduncle on FA skeleton | 25296 |
| Mean L3 in pontine crossing tract on FA skeleton | 25297 |
| Mean L3 in genu of corpus callosum on FA skeleton | 25298 |
| Mean L3 in body of corpus callosum on FA skeleton | 25299 |
| Mean L3 in splenium of corpus callosum on FA skeleton | 25300 |
| Mean L3 in fornix on FA skeleton | 25301 |
| Mean L3 in corticospinal tract on FA skeleton right | 25302 |
| Mean L3 in corticospinal tract on FA skeleton left | 25303 |
| Mean L3 in medial lemniscus on FA skeleton right | 25304 |
| Mean L3 in medial lemniscus on FA skeleton left | 25305 |
| Mean L3 in inferior cerebellar peduncle on FA skeleton right | 25306 |
| Mean L3 in inferior cerebellar peduncle on FA skeleton left | 25307 |
| Mean L3 in superior cerebellar peduncle on FA skeleton right | 25308 |
| Mean L3 in superior cerebellar peduncle on FA skeleton left | 25309 |
| Mean L3 in cerebral peduncle on FA skeleton right | 25310 |
| Mean L3 in cerebral peduncle on FA skeleton left | 25311 |
| Mean L3 in anterior limb of internal capsule on FA skeleton right | 25312 |
| Mean L3 in anterior limb of internal capsule on FA skeleton left | 25313 |
| Mean L3 in posterior limb of internal capsule on FA skeleton right | 25314 |
| Mean L3 in posterior limb of internal capsule on FA skeleton left | 25315 |
| Mean L3 in retrolenticular part of internal capsule on FA skeleton right | 25316 |
| Mean L3 in retrolenticular part of internal capsule on FA skeleton left | 25317 |
| Mean L3 in anterior corona radiata on FA skeleton right | 25318 |
| Mean L3 in anterior corona radiata on FA skeleton left | 25319 |
| Mean L3 in superior corona radiata on FA skeleton right | 25320 |
| Mean L3 in superior corona radiata on FA skeleton left | 25321 |
| Mean L3 in posterior corona radiata on FA skeleton right | 25322 |
| Mean L3 in posterior corona radiata on FA skeleton left | 25323 |
| Mean L3 in posterior thalamic radiation on FA skeleton right | 25324 |
| Mean L3 in posterior thalamic radiation on FA skeleton left | 25325 |
| Mean L3 in sagittal stratum on FA skeleton right | 25326 |
| Mean L3 in sagittal stratum on FA skeleton left | 25327 |
| Mean L3 in external capsule on FA skeleton right | 25328 |
| Mean L3 in external capsule on FA skeleton left | 25329 |
| Mean L3 in cingulum cingulate gyrus on FA skeleton right | 25330 |
| Mean L3 in cingulum cingulate gyrus on FA skeleton left | 25331 |
| Mean L3 in cingulum hippocampus on FA skeleton right | 25332 |
| Mean L3 in cingulum hippocampus on FA skeleton left | 25333 |
| Mean L3 in fornix cres/stria terminalis on FA skeleton right | 25334 |
| Mean L3 in fornix cres/stria terminalis on FA skeleton left | 25335 |
| Mean L3 in superior longitudinal fasciculus on FA skeleton right | 25336 |
| Mean L3 in superior longitudinal fasciculus on FA skeleton left | 25337 |
| Mean L3 in superior frontooccipital fasciculus on FA skeleton right | 25338 |
| Mean L3 in superior frontooccipital fasciculus on FA skeleton left | 25339 |
| Mean L3 in uncinate fasciculus on FA skeleton right | 25340 |
| Mean L3 in uncinate fasciculus on FA skeleton left | 25341 |
| Mean L3 in tapetum on FA skeleton right | 25342 |
| Mean L3 in tapetum on FA skeleton left | 25343 |
| Mean ICVF in middle cerebellar peduncle on FA skeleton | 25344 |
| Mean ICVF in pontine crossing tract on FA skeleton | 25345 |
| Mean ICVF in genu of corpus callosum on FA skeleton | 25346 |
| Mean ICVF in body of corpus callosum on FA skeleton | 25347 |
| Mean ICVF in splenium of corpus callosum on FA skeleton | 25348 |
| Mean ICVF in fornix on FA skeleton | 25349 |
| Mean ICVF in corticospinal tract on FA skeleton right | 25350 |
| Mean ICVF in corticospinal tract on FA skeleton left | 25351 |
| Mean ICVF in medial lemniscus on FA skeleton right | 25352 |
| Mean ICVF in medial lemniscus on FA skeleton left | 25353 |
| Mean ICVF in inferior cerebellar peduncle on FA skeleton right | 25354 |
| Mean ICVF in inferior cerebellar peduncle on FA skeleton left | 25355 |
| Mean ICVF in superior cerebellar peduncle on FA skeleton right | 25356 |
| Mean ICVF in superior cerebellar peduncle on FA skeleton left | 25357 |
| Mean ICVF in cerebral peduncle on FA skeleton right | 25358 |
| Mean ICVF in cerebral peduncle on FA skeleton left | 25359 |
| Mean ICVF in anterior limb of internal capsule on FA skeleton right | 25360 |
| Mean ICVF in anterior limb of internal capsule on FA skeleton left | 25361 |
| Mean ICVF in posterior limb of internal capsule on FA skeleton right | 25362 |
| Mean ICVF in posterior limb of internal capsule on FA skeleton left | 25363 |
| Mean ICVF in retrolenticular part of internal capsule on FA skeleton right | 25364 |
| Mean ICVF in retrolenticular part of internal capsule on FA skeleton left | 25365 |
| Mean ICVF in anterior corona radiata on FA skeleton right | 25366 |
| Mean ICVF in anterior corona radiata on FA skeleton left | 25367 |
| Mean ICVF in superior corona radiata on FA skeleton right | 25368 |
| Mean ICVF in superior corona radiata on FA skeleton left | 25369 |
| Mean ICVF in posterior corona radiata on FA skeleton right | 25370 |
| Mean ICVF in posterior corona radiata on FA skeleton left | 25371 |
| Mean ICVF in posterior thalamic radiation on FA skeleton right | 25372 |
| Mean ICVF in posterior thalamic radiation on FA skeleton left | 25373 |
| Mean ICVF in sagittal stratum on FA skeleton right | 25374 |
| Mean ICVF in sagittal stratum on FA skeleton left | 25375 |
| Mean ICVF in external capsule on FA skeleton right | 25376 |
| Mean ICVF in external capsule on FA skeleton left | 25377 |
| Mean ICVF in cingulum cingulate gyrus on FA skeleton right | 25378 |
| Mean ICVF in cingulum cingulate gyrus on FA skeleton left | 25379 |
| Mean ICVF in cingulum hippocampus on FA skeleton right | 25380 |
| Mean ICVF in cingulum hippocampus on FA skeleton left | 25381 |
| Mean ICVF in fornix cres/stria terminalis on FA skeleton right | 25382 |
| Mean ICVF in fornix cres/stria terminalis on FA skeleton left | 25383 |
| Mean ICVF in superior longitudinal fasciculus on FA skeleton right | 25384 |
| Mean ICVF in superior longitudinal fasciculus on FA skeleton left | 25385 |
| Mean ICVF in superior frontooccipital fasciculus on FA skeleton right | 25386 |
| Mean ICVF in superior frontooccipital fasciculus on FA skeleton left | 25387 |
| Mean ICVF in uncinate fasciculus on FA skeleton right | 25388 |
| Mean ICVF in uncinate fasciculus on FA skeleton left | 25389 |
| Mean ICVF in tapetum on FA skeleton right | 25390 |
| Mean ICVF in tapetum on FA skeleton left | 25391 |
| Mean OD in middle cerebellar peduncle on FA skeleton | 25392 |
| Mean OD in pontine crossing tract on FA skeleton | 25393 |
| Mean OD in genu of corpus callosum on FA skeleton | 25394 |
| Mean OD in body of corpus callosum on FA skeleton | 25395 |
| Mean OD in splenium of corpus callosum on FA skeleton | 25396 |
| Mean OD in fornix on FA skeleton | 25397 |
| Mean OD in corticospinal tract on FA skeleton right | 25398 |
| Mean OD in corticospinal tract on FA skeleton left | 25399 |
| Mean OD in medial lemniscus on FA skeleton right | 25400 |
| Mean OD in medial lemniscus on FA skeleton left | 25401 |
| Mean OD in inferior cerebellar peduncle on FA skeleton right | 25402 |
| Mean OD in inferior cerebellar peduncle on FA skeleton left | 25403 |
| Mean OD in superior cerebellar peduncle on FA skeleton right | 25404 |
| Mean OD in superior cerebellar peduncle on FA skeleton left | 25405 |
| Mean OD in cerebral peduncle on FA skeleton right | 25406 |
| Mean OD in cerebral peduncle on FA skeleton left | 25407 |
| Mean OD in anterior limb of internal capsule on FA skeleton right | 25408 |
| Mean OD in anterior limb of internal capsule on FA skeleton left | 25409 |
| Mean OD in posterior limb of internal capsule on FA skeleton right | 25410 |
| Mean OD in posterior limb of internal capsule on FA skeleton left | 25411 |
| Mean OD in retrolenticular part of internal capsule on FA skeleton right | 25412 |
| Mean OD in retrolenticular part of internal capsule on FA skeleton left | 25413 |
| Mean OD in anterior corona radiata on FA skeleton right | 25414 |
| Mean OD in anterior corona radiata on FA skeleton left | 25415 |
| Mean OD in superior corona radiata on FA skeleton right | 25416 |
| Mean OD in superior corona radiata on FA skeleton left | 25417 |
| Mean OD in posterior corona radiata on FA skeleton right | 25418 |
| Mean OD in posterior corona radiata on FA skeleton left | 25419 |
| Mean OD in posterior thalamic radiation on FA skeleton right | 25420 |
| Mean OD in posterior thalamic radiation on FA skeleton left | 25421 |
| Mean OD in sagittal stratum on FA skeleton right | 25422 |
| Mean OD in sagittal stratum on FA skeleton left | 25423 |
| Mean OD in external capsule on FA skeleton right | 25424 |
| Mean OD in external capsule on FA skeleton left | 25425 |
| Mean OD in cingulum cingulate gyrus on FA skeleton right | 25426 |
| Mean OD in cingulum cingulate gyrus on FA skeleton left | 25427 |
| Mean OD in cingulum hippocampus on FA skeleton right | 25428 |
| Mean OD in cingulum hippocampus on FA skeleton left | 25429 |
| Mean OD in fornix cres/stria terminalis on FA skeleton right | 25430 |
| Mean OD in fornix cres/stria terminalis on FA skeleton left | 25431 |
| Mean OD in superior longitudinal fasciculus on FA skeleton right | 25432 |
| Mean OD in superior longitudinal fasciculus on FA skeleton left | 25433 |
| Mean OD in superior frontooccipital fasciculus on FA skeleton right | 25434 |
| Mean OD in superior frontooccipital fasciculus on FA skeleton left | 25435 |
| Mean OD in uncinate fasciculus on FA skeleton right | 25436 |
| Mean OD in uncinate fasciculus on FA skeleton left | 25437 |
| Mean OD in tapetum on FA skeleton right | 25438 |
| Mean OD in tapetum on FA skeleton left | 25439 |
| Mean ISOVF in middle cerebellar peduncle on FA skeleton | 25440 |
| Mean ISOVF in pontine crossing tract on FA skeleton | 25441 |
| Mean ISOVF in genu of corpus callosum on FA skeleton | 25442 |
| Mean ISOVF in body of corpus callosum on FA skeleton | 25443 |
| Mean ISOVF in splenium of corpus callosum on FA skeleton | 25444 |
| Mean ISOVF in fornix on FA skeleton | 25445 |
| Mean ISOVF in corticospinal tract on FA skeleton right | 25446 |
| Mean ISOVF in corticospinal tract on FA skeleton left | 25447 |
| Mean ISOVF in medial lemniscus on FA skeleton right | 25448 |
| Mean ISOVF in medial lemniscus on FA skeleton left | 25449 |
| Mean ISOVF in inferior cerebellar peduncle on FA skeleton right | 25450 |
| Mean ISOVF in inferior cerebellar peduncle on FA skeleton left | 25451 |
| Mean ISOVF in superior cerebellar peduncle on FA skeleton right | 25452 |
| Mean ISOVF in superior cerebellar peduncle on FA skeleton left | 25453 |
| Mean ISOVF in cerebral peduncle on FA skeleton right | 25454 |
| Mean ISOVF in cerebral peduncle on FA skeleton left | 25455 |
| Mean ISOVF in anterior limb of internal capsule on FA skeleton right | 25456 |
| Mean ISOVF in anterior limb of internal capsule on FA skeleton left | 25457 |
| Mean ISOVF in posterior limb of internal capsule on FA skeleton right | 25458 |
| Mean ISOVF in posterior limb of internal capsule on FA skeleton left | 25459 |
| Mean ISOVF in retrolenticular part of internal capsule on FA skeleton right | 25460 |
| Mean ISOVF in retrolenticular part of internal capsule on FA skeleton left | 25461 |
| Mean ISOVF in anterior corona radiata on FA skeleton right | 25462 |
| Mean ISOVF in anterior corona radiata on FA skeleton left | 25463 |
| Mean ISOVF in superior corona radiata on FA skeleton right | 25464 |
| Mean ISOVF in superior corona radiata on FA skeleton left | 25465 |
| Mean ISOVF in posterior corona radiata on FA skeleton right | 25466 |
| Mean ISOVF in posterior corona radiata on FA skeleton left | 25467 |
| Mean ISOVF in posterior thalamic radiation on FA skeleton right | 25468 |
| Mean ISOVF in posterior thalamic radiation on FA skeleton left | 25469 |
| Mean ISOVF in sagittal stratum on FA skeleton right | 25470 |
| Mean ISOVF in sagittal stratum on FA skeleton left | 25471 |
| Mean ISOVF in external capsule on FA skeleton right | 25472 |
| Mean ISOVF in external capsule on FA skeleton left | 25473 |
| Mean ISOVF in cingulum cingulate gyrus on FA skeleton right | 25474 |
| Mean ISOVF in cingulum cingulate gyrus on FA skeleton left | 25475 |
| Mean ISOVF in cingulum hippocampus on FA skeleton right | 25476 |
| Mean ISOVF in cingulum hippocampus on FA skeleton left | 25477 |
| Mean ISOVF in fornix cres/stria terminalis on FA skeleton right | 25478 |
| Mean ISOVF in fornix cres/stria terminalis on FA skeleton left | 25479 |
| Mean ISOVF in superior longitudinal fasciculus on FA skeleton right | 25480 |
| Mean ISOVF in superior longitudinal fasciculus on FA skeleton left | 25481 |
| Mean ISOVF in superior frontooccipital fasciculus on FA skeleton right | 25482 |
| Mean ISOVF in superior frontooccipital fasciculus on FA skeleton left | 25483 |
| Mean ISOVF in uncinate fasciculus on FA skeleton right | 25484 |
| Mean ISOVF in uncinate fasciculus on FA skeleton left | 25485 |
| Mean ISOVF in tapetum on FA skeleton right | 25486 |
| Mean ISOVF in tapetum on FA skeleton left | 25487 |
| Weighted mean FA in tract acoustic radiation left | 25488 |
| Weighted mean FA in tract acoustic radiation right | 25489 |
| Weighted mean FA in tract anterior thalamic radiation left | 25490 |
| Weighted mean FA in tract anterior thalamic radiation right | 25491 |
| Weighted mean FA in tract cingulate gyrus part of cingulum left | 25492 |
| Weighted mean FA in tract cingulate gyrus part of cingulum right | 25493 |
| Weighted mean FA in tract parahippocampal part of cingulum left | 25494 |
| Weighted mean FA in tract parahippocampal part of cingulum right | 25495 |
| Weighted mean FA in tract corticospinal tract left | 25496 |
| Weighted mean FA in tract corticospinal tract right | 25497 |
| Weighted mean FA in tract forceps major | 25498 |
| Weighted mean FA in tract forceps minor | 25499 |
| Weighted mean FA in tract inferior frontooccipital fasciculus left | 25500 |
| Weighted mean FA in tract inferior frontooccipital fasciculus right | 25501 |
| Weighted mean FA in tract inferior longitudinal fasciculus left | 25502 |
| Weighted mean FA in tract inferior longitudinal fasciculus right | 25503 |
| Weighted mean FA in tract middle cerebellar peduncle | 25504 |
| Weighted mean FA in tract medial lemniscus left | 25505 |
| Weighted mean FA in tract medial lemniscus right | 25506 |
| Weighted mean FA in tract posterior thalamic radiation left | 25507 |
| Weighted mean FA in tract posterior thalamic radiation right | 25508 |
| Weighted mean FA in tract superior longitudinal fasciculus left | 25509 |
| Weighted mean FA in tract superior longitudinal fasciculus right | 25510 |
| Weighted mean FA in tract superior thalamic radiation left | 25511 |
| Weighted mean FA in tract superior thalamic radiation right | 25512 |
| Weighted mean FA in tract uncinate fasciculus left | 25513 |
| Weighted mean FA in tract uncinate fasciculus right | 25514 |
| Weighted mean MD in tract acoustic radiation left | 25515 |
| Weighted mean MD in tract acoustic radiation right | 25516 |
| Weighted mean MD in tract anterior thalamic radiation left | 25517 |
| Weighted mean MD in tract anterior thalamic radiation right | 25518 |
| Weighted mean MD in tract cingulate gyrus part of cingulum left | 25519 |
| Weighted mean MD in tract cingulate gyrus part of cingulum right | 25520 |
| Weighted mean MD in tract parahippocampal part of cingulum left | 25521 |
| Weighted mean MD in tract parahippocampal part of cingulum right | 25522 |
| Weighted mean MD in tract corticospinal tract left | 25523 |
| Weighted mean MD in tract corticospinal tract right | 25524 |
| Weighted mean MD in tract forceps major | 25525 |
| Weighted mean MD in tract forceps minor | 25526 |
| Weighted mean MD in tract inferior frontooccipital fasciculus left | 25527 |
| Weighted mean MD in tract inferior frontooccipital fasciculus right | 25528 |
| Weighted mean MD in tract inferior longitudinal fasciculus left | 25529 |
| Weighted mean MD in tract inferior longitudinal fasciculus right | 25530 |
| Weighted mean MD in tract middle cerebellar peduncle | 25531 |
| Weighted mean MD in tract medial lemniscus left | 25532 |
| Weighted mean MD in tract medial lemniscus right | 25533 |
| Weighted mean MD in tract posterior thalamic radiation left | 25534 |
| Weighted mean MD in tract posterior thalamic radiation right | 25535 |
| Weighted mean MD in tract superior longitudinal fasciculus left | 25536 |
| Weighted mean MD in tract superior longitudinal fasciculus right | 25537 |
| Weighted mean MD in tract superior thalamic radiation left | 25538 |
| Weighted mean MD in tract superior thalamic radiation right | 25539 |
| Weighted mean MD in tract uncinate fasciculus left | 25540 |
| Weighted mean MD in tract uncinate fasciculus right | 25541 |
| Weighted mean MO in tract acoustic radiation left | 25542 |
| Weighted mean MO in tract acoustic radiation right | 25543 |
| Weighted mean MO in tract anterior thalamic radiation left | 25544 |
| Weighted mean MO in tract anterior thalamic radiation right | 25545 |
| Weighted mean MO in tract cingulate gyrus part of cingulum left | 25546 |
| Weighted mean MO in tract cingulate gyrus part of cingulum right | 25547 |
| Weighted mean MO in tract parahippocampal part of cingulum left | 25548 |
| Weighted mean MO in tract parahippocampal part of cingulum right | 25549 |
| Weighted mean MO in tract corticospinal tract left | 25550 |
| Weighted mean MO in tract corticospinal tract right | 25551 |
| Weighted mean MO in tract forceps major | 25552 |
| Weighted mean MO in tract forceps minor | 25553 |
| Weighted mean MO in tract inferior frontooccipital fasciculus left | 25554 |
| Weighted mean MO in tract inferior frontooccipital fasciculus right | 25555 |
| Weighted mean MO in tract inferior longitudinal fasciculus left | 25556 |
| Weighted mean MO in tract inferior longitudinal fasciculus right | 25557 |
| Weighted mean MO in tract middle cerebellar peduncle | 25558 |
| Weighted mean MO in tract medial lemniscus left | 25559 |
| Weighted mean MO in tract medial lemniscus right | 25560 |
| Weighted mean MO in tract posterior thalamic radiation left | 25561 |
| Weighted mean MO in tract posterior thalamic radiation right | 25562 |
| Weighted mean MO in tract superior longitudinal fasciculus left | 25563 |
| Weighted mean MO in tract superior longitudinal fasciculus right | 25564 |
| Weighted mean MO in tract superior thalamic radiation left | 25565 |
| Weighted mean MO in tract superior thalamic radiation right | 25566 |
| Weighted mean MO in tract uncinate fasciculus left | 25567 |
| Weighted mean MO in tract uncinate fasciculus right | 25568 |
| Weighted mean L1 in tract acoustic radiation left | 25569 |
| Weighted mean L1 in tract acoustic radiation right | 25570 |
| Weighted mean L1 in tract anterior thalamic radiation left | 25571 |
| Weighted mean L1 in tract anterior thalamic radiation right | 25572 |
| Weighted mean L1 in tract cingulate gyrus part of cingulum left | 25573 |
| Weighted mean L1 in tract cingulate gyrus part of cingulum right | 25574 |
| Weighted mean L1 in tract parahippocampal part of cingulum left | 25575 |
| Weighted mean L1 in tract parahippocampal part of cingulum right | 25576 |
| Weighted mean L1 in tract corticospinal tract left | 25577 |
| Weighted mean L1 in tract corticospinal tract right | 25578 |
| Weighted mean L1 in tract forceps major | 25579 |
| Weighted mean L1 in tract forceps minor | 25580 |
| Weighted mean L1 in tract inferior frontooccipital fasciculus left | 25581 |
| Weighted mean L1 in tract inferior frontooccipital fasciculus right | 25582 |
| Weighted mean L1 in tract inferior longitudinal fasciculus left | 25583 |
| Weighted mean L1 in tract inferior longitudinal fasciculus right | 25584 |
| Weighted mean L1 in tract middle cerebellar peduncle | 25585 |
| Weighted mean L1 in tract medial lemniscus left | 25586 |
| Weighted mean L1 in tract medial lemniscus right | 25587 |
| Weighted mean L1 in tract posterior thalamic radiation left | 25588 |
| Weighted mean L1 in tract posterior thalamic radiation right | 25589 |
| Weighted mean L1 in tract superior longitudinal fasciculus left | 25590 |
| Weighted mean L1 in tract superior longitudinal fasciculus right | 25591 |
| Weighted mean L1 in tract superior thalamic radiation left | 25592 |
| Weighted mean L1 in tract superior thalamic radiation right | 25593 |
| Weighted mean L1 in tract uncinate fasciculus left | 25594 |
| Weighted mean L1 in tract uncinate fasciculus right | 25595 |
| Weighted mean L2 in tract acoustic radiation left | 25596 |
| Weighted mean L2 in tract acoustic radiation right | 25597 |
| Weighted mean L2 in tract anterior thalamic radiation left | 25598 |
| Weighted mean L2 in tract anterior thalamic radiation right | 25599 |
| Weighted mean L2 in tract cingulate gyrus part of cingulum left | 25600 |
| Weighted mean L2 in tract cingulate gyrus part of cingulum right | 25601 |
| Weighted mean L2 in tract parahippocampal part of cingulum left | 25602 |
| Weighted mean L2 in tract parahippocampal part of cingulum right | 25603 |
| Weighted mean L2 in tract corticospinal tract left | 25604 |
| Weighted mean L2 in tract corticospinal tract right | 25605 |
| Weighted mean L2 in tract forceps major | 25606 |
| Weighted mean L2 in tract forceps minor | 25607 |
| Weighted mean L2 in tract inferior frontooccipital fasciculus left | 25608 |
| Weighted mean L2 in tract inferior frontooccipital fasciculus right | 25609 |
| Weighted mean L2 in tract inferior longitudinal fasciculus left | 25610 |
| Weighted mean L2 in tract inferior longitudinal fasciculus right | 25611 |
| Weighted mean L2 in tract middle cerebellar peduncle | 25612 |
| Weighted mean L2 in tract medial lemniscus left | 25613 |
| Weighted mean L2 in tract medial lemniscus right | 25614 |
| Weighted mean L2 in tract posterior thalamic radiation left | 25615 |
| Weighted mean L2 in tract posterior thalamic radiation right | 25616 |
| Weighted mean L2 in tract superior longitudinal fasciculus left | 25617 |
| Weighted mean L2 in tract superior longitudinal fasciculus right | 25618 |
| Weighted mean L2 in tract superior thalamic radiation left | 25619 |
| Weighted mean L2 in tract superior thalamic radiation right | 25620 |
| Weighted mean L2 in tract uncinate fasciculus left | 25621 |
| Weighted mean L2 in tract uncinate fasciculus right | 25622 |
| Weighted mean L3 in tract acoustic radiation left | 25623 |
| Weighted mean L3 in tract acoustic radiation right | 25624 |
| Weighted mean L3 in tract anterior thalamic radiation left | 25625 |
| Weighted mean L3 in tract anterior thalamic radiation right | 25626 |
| Weighted mean L3 in tract cingulate gyrus part of cingulum left | 25627 |
| Weighted mean L3 in tract cingulate gyrus part of cingulum right | 25628 |
| Weighted mean L3 in tract parahippocampal part of cingulum left | 25629 |
| Weighted mean L3 in tract parahippocampal part of cingulum right | 25630 |
| Weighted mean L3 in tract corticospinal tract left | 25631 |
| Weighted mean L3 in tract corticospinal tract right | 25632 |
| Weighted mean L3 in tract forceps major | 25633 |
| Weighted mean L3 in tract forceps minor | 25634 |
| Weighted mean L3 in tract inferior frontooccipital fasciculus left | 25635 |
| Weighted mean L3 in tract inferior frontooccipital fasciculus right | 25636 |
| Weighted mean L3 in tract inferior longitudinal fasciculus left | 25637 |
| Weighted mean L3 in tract inferior longitudinal fasciculus right | 25638 |
| Weighted mean L3 in tract middle cerebellar peduncle | 25639 |
| Weighted mean L3 in tract medial lemniscus left | 25640 |
| Weighted mean L3 in tract medial lemniscus right | 25641 |
| Weighted mean L3 in tract posterior thalamic radiation left | 25642 |
| Weighted mean L3 in tract posterior thalamic radiation right | 25643 |
| Weighted mean L3 in tract superior longitudinal fasciculus left | 25644 |
| Weighted mean L3 in tract superior longitudinal fasciculus right | 25645 |
| Weighted mean L3 in tract superior thalamic radiation left | 25646 |
| Weighted mean L3 in tract superior thalamic radiation right | 25647 |
| Weighted mean L3 in tract uncinate fasciculus left | 25648 |
| Weighted mean L3 in tract uncinate fasciculus right | 25649 |
| Weighted mean ICVF in tract acoustic radiation left | 25650 |
| Weighted mean ICVF in tract acoustic radiation right | 25651 |
| Weighted mean ICVF in tract anterior thalamic radiation left | 25652 |
| Weighted mean ICVF in tract anterior thalamic radiation right | 25653 |
| Weighted mean ICVF in tract cingulate gyrus part of cingulum left | 25654 |
| Weighted mean ICVF in tract cingulate gyrus part of cingulum right | 25655 |
| Weighted mean ICVF in tract parahippocampal part of cingulum left | 25656 |
| Weighted mean ICVF in tract parahippocampal part of cingulum right | 25657 |
| Weighted mean ICVF in tract corticospinal tract left | 25658 |
| Weighted mean ICVF in tract corticospinal tract right | 25659 |
| Weighted mean ICVF in tract forceps major | 25660 |
| Weighted mean ICVF in tract forceps minor | 25661 |
| Weighted mean ICVF in tract inferior frontooccipital fasciculus left | 25662 |
| Weighted mean ICVF in tract inferior frontooccipital fasciculus right | 25663 |
| Weighted mean ICVF in tract inferior longitudinal fasciculus left | 25664 |
| Weighted mean ICVF in tract inferior longitudinal fasciculus right | 25665 |
| Weighted mean ICVF in tract middle cerebellar peduncle | 25666 |
| Weighted mean ICVF in tract medial lemniscus left | 25667 |
| Weighted mean ICVF in tract medial lemniscus right | 25668 |
| Weighted mean ICVF in tract posterior thalamic radiation left | 25669 |
| Weighted mean ICVF in tract posterior thalamic radiation right | 25670 |
| Weighted mean ICVF in tract superior longitudinal fasciculus left | 25671 |
| Weighted mean ICVF in tract superior longitudinal fasciculus right | 25672 |
| Weighted mean ICVF in tract superior thalamic radiation left | 25673 |
| Weighted mean ICVF in tract superior thalamic radiation right | 25674 |
| Weighted mean ICVF in tract uncinate fasciculus left | 25675 |
| Weighted mean ICVF in tract uncinate fasciculus right | 25676 |
| Weighted mean OD in tract acoustic radiation left | 25677 |
| Weighted mean OD in tract acoustic radiation right | 25678 |
| Weighted mean OD in tract anterior thalamic radiation left | 25679 |
| Weighted mean OD in tract anterior thalamic radiation right | 25680 |
| Weighted mean OD in tract cingulate gyrus part of cingulum left | 25681 |
| Weighted mean OD in tract cingulate gyrus part of cingulum right | 25682 |
| Weighted mean OD in tract parahippocampal part of cingulum left | 25683 |
| Weighted mean OD in tract parahippocampal part of cingulum right | 25684 |
| Weighted mean OD in tract corticospinal tract left | 25685 |
| Weighted mean OD in tract corticospinal tract right | 25686 |
| Weighted mean OD in tract forceps major | 25687 |
| Weighted mean OD in tract forceps minor | 25688 |
| Weighted mean OD in tract inferior frontooccipital fasciculus left | 25689 |
| Weighted mean OD in tract inferior frontooccipital fasciculus right | 25690 |
| Weighted mean OD in tract inferior longitudinal fasciculus left | 25691 |
| Weighted mean OD in tract inferior longitudinal fasciculus right | 25692 |
| Weighted mean OD in tract middle cerebellar peduncle | 25693 |
| Weighted mean OD in tract medial lemniscus left | 25694 |
| Weighted mean OD in tract medial lemniscus right | 25695 |
| Weighted mean OD in tract posterior thalamic radiation left | 25696 |
| Weighted mean OD in tract posterior thalamic radiation right | 25697 |
| Weighted mean OD in tract superior longitudinal fasciculus left | 25698 |
| Weighted mean OD in tract superior longitudinal fasciculus right | 25699 |
| Weighted mean OD in tract superior thalamic radiation left | 25700 |
| Weighted mean OD in tract superior thalamic radiation right | 25701 |
| Weighted mean OD in tract uncinate fasciculus left | 25702 |
| Weighted mean OD in tract uncinate fasciculus right | 25703 |
| Weighted mean ISOVF in tract acoustic radiation left | 25704 |
| Weighted mean ISOVF in tract acoustic radiation right | 25705 |
| Weighted mean ISOVF in tract anterior thalamic radiation left | 25706 |
| Weighted mean ISOVF in tract anterior thalamic radiation right | 25707 |
| Weighted mean ISOVF in tract cingulate gyrus part of cingulum left | 25708 |
| Weighted mean ISOVF in tract cingulate gyrus part of cingulum right | 25709 |
| Weighted mean ISOVF in tract parahippocampal part of cingulum left | 25710 |
| Weighted mean ISOVF in tract parahippocampal part of cingulum right | 25711 |
| Weighted mean ISOVF in tract corticospinal tract left | 25712 |
| Weighted mean ISOVF in tract corticospinal tract right | 25713 |
| Weighted mean ISOVF in tract forceps major | 25714 |
| Weighted mean ISOVF in tract forceps minor | 25715 |
| Weighted mean ISOVF in tract inferior frontooccipital fasciculus left | 25716 |
| Weighted mean ISOVF in tract inferior frontooccipital fasciculus right | 25717 |
| Weighted mean ISOVF in tract inferior longitudinal fasciculus left | 25718 |
| Weighted mean ISOVF in tract inferior longitudinal fasciculus right | 25719 |
| Weighted mean ISOVF in tract middle cerebellar peduncle | 25720 |
| Weighted mean ISOVF in tract medial lemniscus left | 25721 |
| Weighted mean ISOVF in tract medial lemniscus right | 25722 |
| Weighted mean ISOVF in tract posterior thalamic radiation left | 25723 |
| Weighted mean ISOVF in tract posterior thalamic radiation right | 25724 |
| Weighted mean ISOVF in tract superior longitudinal fasciculus left | 25725 |
| Weighted mean ISOVF in tract superior longitudinal fasciculus right | 25726 |
| Weighted mean ISOVF in tract superior thalamic radiation left | 25727 |
| Weighted mean ISOVF in tract superior thalamic radiation right | 25728 |
| Weighted mean ISOVF in tract uncinate fasciculus left | 25729 |
| Weighted mean ISOVF in tract uncinate fasciculus right | 25730 |
| Resting-state partial correlation 25-dimension IC v001 | 25752 |
| Resting-state partial correlation 25-dimension IC v002 | 25752 |
| Resting-state partial correlation 25-dimension IC v003 | 25752 |
| Resting-state partial correlation 25-dimension IC v004 | 25752 |
| Resting-state partial correlation 25-dimension IC v005 | 25752 |
| Resting-state partial correlation 25-dimension IC v006 | 25752 |
| Resting-state partial correlation 25-dimension IC v007 | 25752 |
| Resting-state partial correlation 25-dimension IC v008 | 25752 |
| Resting-state partial correlation 25-dimension IC v009 | 25752 |
| Resting-state partial correlation 25-dimension IC v010 | 25752 |
| Resting-state partial correlation 25-dimension IC v011 | 25752 |
| Resting-state partial correlation 25-dimension IC v012 | 25752 |
| Resting-state partial correlation 25-dimension IC v013 | 25752 |
| Resting-state partial correlation 25-dimension IC v014 | 25752 |
| Resting-state partial correlation 25-dimension IC v015 | 25752 |
| Resting-state partial correlation 25-dimension IC v016 | 25752 |
| Resting-state partial correlation 25-dimension IC v017 | 25752 |
| Resting-state partial correlation 25-dimension IC v018 | 25752 |
| Resting-state partial correlation 25-dimension IC v019 | 25752 |
| Resting-state partial correlation 25-dimension IC v020 | 25752 |
| Resting-state partial correlation 25-dimension IC v021 | 25752 |
| Resting-state partial correlation 25-dimension IC v022 | 25752 |
| Resting-state partial correlation 25-dimension IC v023 | 25752 |
| Resting-state partial correlation 25-dimension IC v024 | 25752 |
| Resting-state partial correlation 25-dimension IC v025 | 25752 |
| Resting-state partial correlation 25-dimension IC v026 | 25752 |
| Resting-state partial correlation 25-dimension IC v027 | 25752 |
| Resting-state partial correlation 25-dimension IC v028 | 25752 |
| Resting-state partial correlation 25-dimension IC v029 | 25752 |
| Resting-state partial correlation 25-dimension IC v030 | 25752 |
| Resting-state partial correlation 25-dimension IC v031 | 25752 |
| Resting-state partial correlation 25-dimension IC v032 | 25752 |
| Resting-state partial correlation 25-dimension IC v033 | 25752 |
| Resting-state partial correlation 25-dimension IC v034 | 25752 |
| Resting-state partial correlation 25-dimension IC v035 | 25752 |
| Resting-state partial correlation 25-dimension IC v036 | 25752 |
| Resting-state partial correlation 25-dimension IC v037 | 25752 |
| Resting-state partial correlation 25-dimension IC v038 | 25752 |
| Resting-state partial correlation 25-dimension IC v039 | 25752 |
| Resting-state partial correlation 25-dimension IC v040 | 25752 |
| Resting-state partial correlation 25-dimension IC v041 | 25752 |
| Resting-state partial correlation 25-dimension IC v042 | 25752 |
| Resting-state partial correlation 25-dimension IC v043 | 25752 |
| Resting-state partial correlation 25-dimension IC v044 | 25752 |
| Resting-state partial correlation 25-dimension IC v045 | 25752 |
| Resting-state partial correlation 25-dimension IC v046 | 25752 |
| Resting-state partial correlation 25-dimension IC v047 | 25752 |
| Resting-state partial correlation 25-dimension IC v048 | 25752 |
| Resting-state partial correlation 25-dimension IC v049 | 25752 |
| Resting-state partial correlation 25-dimension IC v050 | 25752 |
| Resting-state partial correlation 25-dimension IC v051 | 25752 |
| Resting-state partial correlation 25-dimension IC v052 | 25752 |
| Resting-state partial correlation 25-dimension IC v053 | 25752 |
| Resting-state partial correlation 25-dimension IC v054 | 25752 |
| Resting-state partial correlation 25-dimension IC v055 | 25752 |
| Resting-state partial correlation 25-dimension IC v056 | 25752 |
| Resting-state partial correlation 25-dimension IC v057 | 25752 |
| Resting-state partial correlation 25-dimension IC v058 | 25752 |
| Resting-state partial correlation 25-dimension IC v059 | 25752 |
| Resting-state partial correlation 25-dimension IC v060 | 25752 |
| Resting-state partial correlation 25-dimension IC v061 | 25752 |
| Resting-state partial correlation 25-dimension IC v062 | 25752 |
| Resting-state partial correlation 25-dimension IC v063 | 25752 |
| Resting-state partial correlation 25-dimension IC v064 | 25752 |
| Resting-state partial correlation 25-dimension IC v065 | 25752 |
| Resting-state partial correlation 25-dimension IC v066 | 25752 |
| Resting-state partial correlation 25-dimension IC v067 | 25752 |
| Resting-state partial correlation 25-dimension IC v068 | 25752 |
| Resting-state partial correlation 25-dimension IC v069 | 25752 |
| Resting-state partial correlation 25-dimension IC v070 | 25752 |
| Resting-state partial correlation 25-dimension IC v071 | 25752 |
| Resting-state partial correlation 25-dimension IC v072 | 25752 |
| Resting-state partial correlation 25-dimension IC v073 | 25752 |
| Resting-state partial correlation 25-dimension IC v074 | 25752 |
| Resting-state partial correlation 25-dimension IC v075 | 25752 |
| Resting-state partial correlation 25-dimension IC v076 | 25752 |
| Resting-state partial correlation 25-dimension IC v077 | 25752 |
| Resting-state partial correlation 25-dimension IC v078 | 25752 |
| Resting-state partial correlation 25-dimension IC v079 | 25752 |
| Resting-state partial correlation 25-dimension IC v080 | 25752 |
| Resting-state partial correlation 25-dimension IC v081 | 25752 |
| Resting-state partial correlation 25-dimension IC v082 | 25752 |
| Resting-state partial correlation 25-dimension IC v083 | 25752 |
| Resting-state partial correlation 25-dimension IC v084 | 25752 |
| Resting-state partial correlation 25-dimension IC v085 | 25752 |
| Resting-state partial correlation 25-dimension IC v086 | 25752 |
| Resting-state partial correlation 25-dimension IC v087 | 25752 |
| Resting-state partial correlation 25-dimension IC v088 | 25752 |
| Resting-state partial correlation 25-dimension IC v089 | 25752 |
| Resting-state partial correlation 25-dimension IC v090 | 25752 |
| Resting-state partial correlation 25-dimension IC v091 | 25752 |
| Resting-state partial correlation 25-dimension IC v092 | 25752 |
| Resting-state partial correlation 25-dimension IC v093 | 25752 |
| Resting-state partial correlation 25-dimension IC v094 | 25752 |
| Resting-state partial correlation 25-dimension IC v095 | 25752 |
| Resting-state partial correlation 25-dimension IC v096 | 25752 |
| Resting-state partial correlation 25-dimension IC v097 | 25752 |
| Resting-state partial correlation 25-dimension IC v098 | 25752 |
| Resting-state partial correlation 25-dimension IC v099 | 25752 |
| Resting-state partial correlation 25-dimension IC v100 | 25752 |
| Resting-state partial correlation 25-dimension IC v101 | 25752 |
| Resting-state partial correlation 25-dimension IC v102 | 25752 |
| Resting-state partial correlation 25-dimension IC v103 | 25752 |
| Resting-state partial correlation 25-dimension IC v104 | 25752 |
| Resting-state partial correlation 25-dimension IC v105 | 25752 |
| Resting-state partial correlation 25-dimension IC v106 | 25752 |
| Resting-state partial correlation 25-dimension IC v107 | 25752 |
| Resting-state partial correlation 25-dimension IC v108 | 25752 |
| Resting-state partial correlation 25-dimension IC v109 | 25752 |
| Resting-state partial correlation 25-dimension IC v110 | 25752 |
| Resting-state partial correlation 25-dimension IC v111 | 25752 |
| Resting-state partial correlation 25-dimension IC v112 | 25752 |
| Resting-state partial correlation 25-dimension IC v113 | 25752 |
| Resting-state partial correlation 25-dimension IC v114 | 25752 |
| Resting-state partial correlation 25-dimension IC v115 | 25752 |
| Resting-state partial correlation 25-dimension IC v116 | 25752 |
| Resting-state partial correlation 25-dimension IC v117 | 25752 |
| Resting-state partial correlation 25-dimension IC v118 | 25752 |
| Resting-state partial correlation 25-dimension IC v119 | 25752 |
| Resting-state partial correlation 25-dimension IC v120 | 25752 |
| Resting-state partial correlation 25-dimension IC v121 | 25752 |
| Resting-state partial correlation 25-dimension IC v122 | 25752 |
| Resting-state partial correlation 25-dimension IC v123 | 25752 |
| Resting-state partial correlation 25-dimension IC v124 | 25752 |
| Resting-state partial correlation 25-dimension IC v125 | 25752 |
| Resting-state partial correlation 25-dimension IC v126 | 25752 |
| Resting-state partial correlation 25-dimension IC v127 | 25752 |
| Resting-state partial correlation 25-dimension IC v128 | 25752 |
| Resting-state partial correlation 25-dimension IC v129 | 25752 |
| Resting-state partial correlation 25-dimension IC v130 | 25752 |
| Resting-state partial correlation 25-dimension IC v131 | 25752 |
| Resting-state partial correlation 25-dimension IC v132 | 25752 |
| Resting-state partial correlation 25-dimension IC v133 | 25752 |
| Resting-state partial correlation 25-dimension IC v134 | 25752 |
| Resting-state partial correlation 25-dimension IC v135 | 25752 |
| Resting-state partial correlation 25-dimension IC v136 | 25752 |
| Resting-state partial correlation 25-dimension IC v137 | 25752 |
| Resting-state partial correlation 25-dimension IC v138 | 25752 |
| Resting-state partial correlation 25-dimension IC v139 | 25752 |
| Resting-state partial correlation 25-dimension IC v140 | 25752 |
| Resting-state partial correlation 25-dimension IC v141 | 25752 |
| Resting-state partial correlation 25-dimension IC v142 | 25752 |
| Resting-state partial correlation 25-dimension IC v143 | 25752 |
| Resting-state partial correlation 25-dimension IC v144 | 25752 |
| Resting-state partial correlation 25-dimension IC v145 | 25752 |
| Resting-state partial correlation 25-dimension IC v146 | 25752 |
| Resting-state partial correlation 25-dimension IC v147 | 25752 |
| Resting-state partial correlation 25-dimension IC v148 | 25752 |
| Resting-state partial correlation 25-dimension IC v149 | 25752 |
| Resting-state partial correlation 25-dimension IC v150 | 25752 |
| Resting-state partial correlation 25-dimension IC v151 | 25752 |
| Resting-state partial correlation 25-dimension IC v152 | 25752 |
| Resting-state partial correlation 25-dimension IC v153 | 25752 |
| Resting-state partial correlation 25-dimension IC v154 | 25752 |
| Resting-state partial correlation 25-dimension IC v155 | 25752 |
| Resting-state partial correlation 25-dimension IC v156 | 25752 |
| Resting-state partial correlation 25-dimension IC v157 | 25752 |
| Resting-state partial correlation 25-dimension IC v158 | 25752 |
| Resting-state partial correlation 25-dimension IC v159 | 25752 |
| Resting-state partial correlation 25-dimension IC v160 | 25752 |
| Resting-state partial correlation 25-dimension IC v161 | 25752 |
| Resting-state partial correlation 25-dimension IC v162 | 25752 |
| Resting-state partial correlation 25-dimension IC v163 | 25752 |
| Resting-state partial correlation 25-dimension IC v164 | 25752 |
| Resting-state partial correlation 25-dimension IC v165 | 25752 |
| Resting-state partial correlation 25-dimension IC v166 | 25752 |
| Resting-state partial correlation 25-dimension IC v167 | 25752 |
| Resting-state partial correlation 25-dimension IC v168 | 25752 |
| Resting-state partial correlation 25-dimension IC v169 | 25752 |
| Resting-state partial correlation 25-dimension IC v170 | 25752 |
| Resting-state partial correlation 25-dimension IC v171 | 25752 |
| Resting-state partial correlation 25-dimension IC v172 | 25752 |
| Resting-state partial correlation 25-dimension IC v173 | 25752 |
| Resting-state partial correlation 25-dimension IC v174 | 25752 |
| Resting-state partial correlation 25-dimension IC v175 | 25752 |
| Resting-state partial correlation 25-dimension IC v176 | 25752 |
| Resting-state partial correlation 25-dimension IC v177 | 25752 |
| Resting-state partial correlation 25-dimension IC v178 | 25752 |
| Resting-state partial correlation 25-dimension IC v179 | 25752 |
| Resting-state partial correlation 25-dimension IC v180 | 25752 |
| Resting-state partial correlation 25-dimension IC v181 | 25752 |
| Resting-state partial correlation 25-dimension IC v182 | 25752 |
| Resting-state partial correlation 25-dimension IC v183 | 25752 |
| Resting-state partial correlation 25-dimension IC v184 | 25752 |
| Resting-state partial correlation 25-dimension IC v185 | 25752 |
| Resting-state partial correlation 25-dimension IC v186 | 25752 |
| Resting-state partial correlation 25-dimension IC v187 | 25752 |
| Resting-state partial correlation 25-dimension IC v188 | 25752 |
| Resting-state partial correlation 25-dimension IC v189 | 25752 |
| Resting-state partial correlation 25-dimension IC v190 | 25752 |
| Resting-state partial correlation 25-dimension IC v191 | 25752 |
| Resting-state partial correlation 25-dimension IC v192 | 25752 |
| Resting-state partial correlation 25-dimension IC v193 | 25752 |
| Resting-state partial correlation 25-dimension IC v194 | 25752 |
| Resting-state partial correlation 25-dimension IC v195 | 25752 |
| Resting-state partial correlation 25-dimension IC v196 | 25752 |
| Resting-state partial correlation 25-dimension IC v197 | 25752 |
| Resting-state partial correlation 25-dimension IC v198 | 25752 |
| Resting-state partial correlation 25-dimension IC v199 | 25752 |
| Resting-state partial correlation 25-dimension IC v200 | 25752 |
| Resting-state partial correlation 25-dimension IC v201 | 25752 |
| Resting-state partial correlation 25-dimension IC v202 | 25752 |
| Resting-state partial correlation 25-dimension IC v203 | 25752 |
| Resting-state partial correlation 25-dimension IC v204 | 25752 |
| Resting-state partial correlation 25-dimension IC v205 | 25752 |
| Resting-state partial correlation 25-dimension IC v206 | 25752 |
| Resting-state partial correlation 25-dimension IC v207 | 25752 |
| Resting-state partial correlation 25-dimension IC v208 | 25752 |
| Resting-state partial correlation 25-dimension IC v209 | 25752 |
| Resting-state partial correlation 25-dimension IC v210 | 25752 |

ICVF = intracellular volume fraction; FA = fractional anisotropy; ISOVF = isotropic volume fraction; MO = mode of anisotropy; OD = orientation dispersion; v# = variable from resting-state fMRI partial correlation matrix, using 25-dimension independent component analysis.
